# Structural Basis for Promoter Recognition and Transcription Factor Binding and Release in Human Mitochondria

**DOI:** 10.1101/2025.04.03.647028

**Authors:** Karl Herbine, Ashok R. Nayak, Angelica Zamudio-Ochoa, Dmitry Temiakov

**Affiliations:** Department of Biochemistry and Molecular Biology, Thomas Jefferson University; 1020 Locust St, Philadelphia 19107, USA

**Keywords:** mitochondrial transcription, POLRMT, mtRNAP, TFAM, TFB2M, TEFM

## Abstract

Transcription in human mitochondria is driven by a core apparatus consisting of a Pol A family RNA polymerase (mtRNAP), the initiation factors TFAM and TFB2M, and the elongation factor TEFM. While earlier structures of initiation and elongation complexes provided valuable snapshots, they represent isolated stages of a highly dynamic and multistep process. Critical aspects of mitochondrial transcription—such as DNA recognition and melting, promoter escape, and the release of initiation factors—remain poorly understood. Here, we present a series of cryo-EM structures that capture the transcription complex as it transitions from the initial open promoter complex to the processive elongation complex through intermediate stages. Our data reveal new determinants of promoter specificity, the sequential disengagement of mtRNAP from TFAM and the promoter, the release of TFB2M, and the recruitment of TEFM. Together, these findings provide a detailed molecular mechanism underlying transcription in human mitochondria.

## INTRODUCTION

Mitochondria, the energy-producing organelles, house a maternally inherited genome that encodes subunits of oxidative phosphorylation complexes. Defects in mitochondrial gene expression lead to severe diseases, including mitochondrial myopathies, neurodegenerative disorders, progressive external ophthalmoplegia, and systemic conditions such as MELAS and Leigh syndrome (Vanderslice et al., 1986; Wallace, 2018). These disorders often stem from impaired transcription, replication, or translation of mitochondrial DNA, resulting in energy deficits and cellular dysfunction (Gustafson et al., 2020). Understanding mitochondrial transcription is crucial for developing targeted therapies, as it represents a promising yet underexplored therapeutic target for mitochondrial and neurodegenerative diseases, and cancer (Bonekamp et al., 2020; Olahova et al., 2021; Van Haute et al., 2023).

A key difference between mitochondrial and nuclear transcription lies in its unique core transcription apparatus (Falkenberg et al., 2002; Fisher and Clayton, 1988; Masters et al., 1987). This apparatus consists of a single-subunit phage-type RNA polymerase (mtRNAP, also known as POLRMT) and a dedicated set of bacterial-origin transcription factors, and is tasked with the synthesis of mRNA, tRNA, rRNA, and replication primers (Hillen et al., 2018; Ringel et al., 2011). Major unresolved questions in mitochondrial transcription include how the transcription machinery regulates mRNA synthesis from the multi-copy mitochondrial genome, aligning it with the cell’s energetic demands, and how it transitions from processive transcription to generating primers necessary for mtDNA replication (Hillen *et al*., 2018; Tan et al., 2023).

Human mtRNAP relies on two transcription initiation factors, TFAM and TFB2M, as well as an elongation factor, TEFM, for processive RNA synthesis (Litonin et al., 2010; Tan *et al*., 2023). TFAM, a major nucleoid protein that protects mtDNA by organizing it into nucleoids, binds to the –40/-17 region of the promoter, inducing its 180° bending (Rubio-Cosials and Sola, 2013). According to the sequential model of mitochondrial transcription initiation (Morozov et al., 2014; Posse et al., 2014), TFAM recruits mtRNAP to the promoter, positioning the polymerase’s active site over the transcription start site and forming a pre-initiation complex. Promoter opening requires the recruitment of another initiation factor, TFB2M, a repurposed (and catalytically inactive) methyltransferase. TFB2M binds to the single-stranded region of the non-template (NT) strand, stabilizing the open promoter complex (Hillen et al., 2017a; Sologub et al., 2009).

During the transition to the elongation phase of transcription, initiation factors are released, and mtRNAP binds TEFM, a non-functional Holliday junction resolvase that has also been repurposed in human mitochondria (Minczuk et al., 2011). TEFM binds at a site that overlaps with TFB2M’s binding region on mtRNAP and is essential for transcription complex stability and processivity (Hillen et al., 2017b). Additionally, TEFM functions as an antiterminator, allowing the polymerase to transcribe through G-quadruplex structures that would otherwise cause premature transcription termination and generate replication primers (Agaronyan et al., 2015; Hillen *et al*., 2017b; Posse et al., 2015). The stage of mitochondrial transcription at which the initiation factors are released and the elongation factor is recruited remains unknown (Hillen *et al*., 2018).

In this study, we employed single-particle cryo-EM to capture the major intermediate states of mitochondrial transcription initiation, providing a detailed structural and mechanistic framework for this process. The obtained high-resolution structures represent the initial binding and recognition of the promoter, promoter melting, initial RNA synthesis, TFAM and TFB2M dissociation, the transition to the elongation phase of transcription, and the recruitment of the elongation factor TEFM. By elucidating these key steps, we provide insights into the dynamics of the mitochondrial transcription machinery and its regulation, contributing to our understanding of how mitochondrial gene expression aligns with cellular energetic demands.

## RESULTS

### Capturing the Early Stages of Mitochondrial Transcription Initiation

Transcription initiation in human mitochondria involves mtRNAP and core initiation factors TFAM and TFB2M (**Figure 1A**), which form a ∼ 200 kDa protein complex with DNA. To investigate the early stages of transcription, we assembled the initiation complex (IC) on promoter DNA containing a 4-nucleotide non-complementary region near the transcription start site (TSS), spanning bases –3 to +1 (**Figure 1B**). Pre-melting of the promoter near the TSS is essential for capturing the open promoter complex (IC0), which, in the absence of substrate NTPs or short RNA primers, is inherently unstable and complicates cryo-EM studies (Morozov *et al*., 2014).

**Figure 1.**
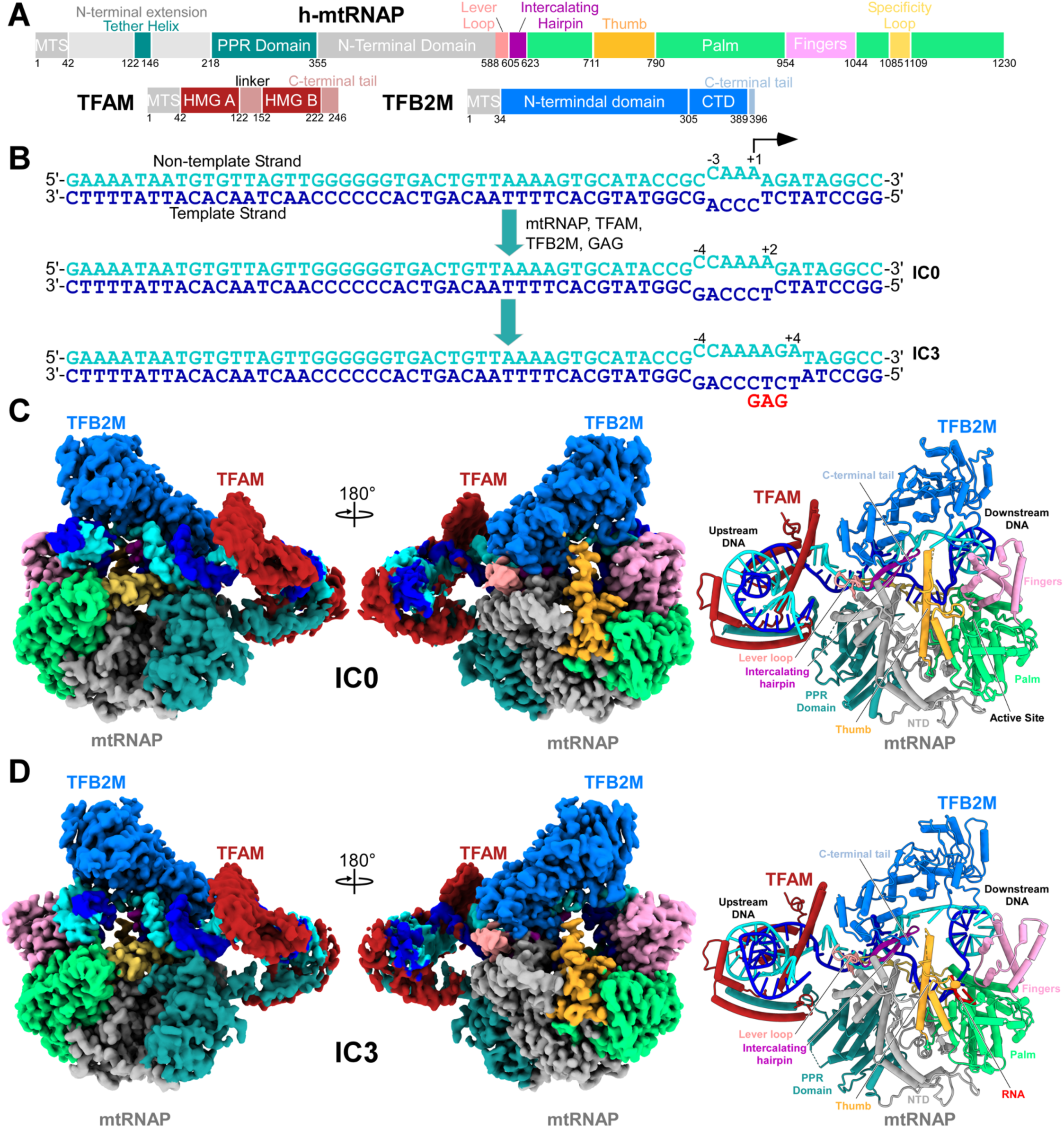
Structure of the Early-Stage Transcription Initiation Complex. **A.** Schematic structure of mtRNAP, TFAM, and TFB2M. The coloring of the major domains and structural elements is shown throughout all figures. **B.** Design of the promoter template. The template strand of DNA (blue) contains a non-complementary region, allowing for a 4 bp pre-melted region. This region expands upon the formation of IC0 and IC3. The non-template strand is shown in cyan. The black arrow at the +1 base indicates the transcription start site. **C.** The cryo-EM density and the structure of IC0. TFB2M is shown in marine, TFAM is dark red. Prominent structural elements of mtRNAP are indicated. **D.** The cryo-EM density and the structure of IC3.

Because our previous studies suggested that TFB2M recognizes the NT strand in the region near the TSS (Hillen *et al*., 2017a; Zamudio-Ochoa et al., 2022), we preserved its sequence and modified the template (T) strand instead (**Figure 1B**). Additionally, to prevent transcript slippage and reduce heterogeneity of the IC, we substituted the templating +1 dTMP base to dCMP and provided a 3-mer RNA primer, GAG, to stabilize the IC (Zamudio-Ochoa *et al*., 2022). Utilization of short 2-3 nt RNA primers is a well-documented phenomenon representing RNA recycling by various RNA polymerases (Goldman et al., 2011).

Initial single-particle analysis (SPA) and 3D reconstruction revealed strong orientation bias and weak TFAM density in ICs, likely caused by the flexibility of its HMG-box domains. Indeed, recent single-molecule fluorescence resonance energy transfer (smFRET) studies have shown that TFAM binding to mtDNA adopts a partially bent, flexible hinge state, which can diffusely slide along the mtDNA (Huh et al., 2024).

SPA optimization sought to address challenges associated with the limited number of particles representing complete ICs by utilizing 3DFlex, a computational method designed to model the flexible and continuous conformational variability of macromolecular structures (Punjani and Fleet, 2023) (**Figure S1**). However, even when combined with local refinement and particle subtraction, this approach resulted in only marginal improvement in the TFAM signal within the density maps, indicating that compositional heterogeneity or the presence of intermediate states may be limiting the resolution of the dataset.

To overcome this, we broadened the particle selection criteria and proceeded directly to *ab initio* 3D reconstruction without intervening 2D-classification steps (**Figure S1, Table S1**). This was followed by multiple rounds of supervised heterogeneous refinement, enabling a more thorough evaluation of the dataset without prematurely discarding potentially informative particles. This strategy allowed capturing rare particle orientations that were originally obscured by prior stringent 2D classification and resulted in enhanced structural detail across the dataset (**Figure S1, S2, S3**).

Two major conformations of the IC were revealed by SPA (**Figure 1C, D, S1**). In Structure ONE, the pre-melted region expanded from 4 to 6 bp, spanning bases –4 to +2, representing the initially open promoter complex (referred to as IC0; **Figure 1C**). In IC0, the T strand had not yet descended into the active site, and no bound RNA product was observed in the complex (**Figure 2A**). Additionally, the fingers domain of mtRNAP adopted a clenched conformation, previously observed in the apo form of mtRNAP (Ringel *et al*., 2011), suggesting that IC0 represents a catalytically inactive state (**Figure 2A**).

**Figure 2.**
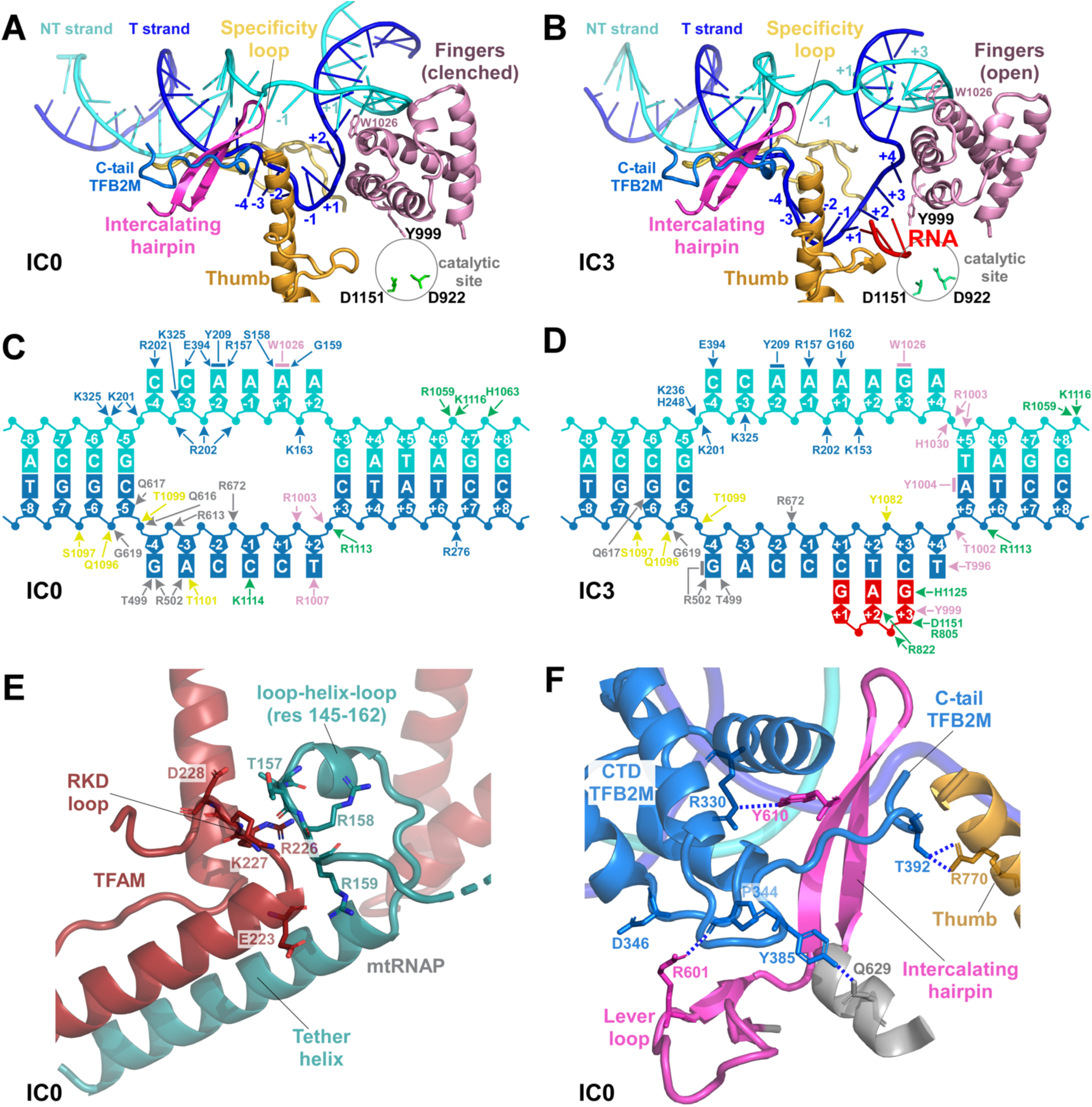
Interactions with the Transcription Initiation Complex. **A,B.** The trajectory of the nucleic acids in IC0 (A) and IC3 (B). Structures are aligned with respect to their conserved palm subdomains. Major structural elements of mtRNAP and the C-tail of TFB2M are indicated. The position of the active site is indicated by a black circle. The catalytic aspartate residues are shown as sticks. **C,D.** A network of key interactions between the nucleic acids and proteins in IC0 (**C**) and IC3 (**D**). Arrows indicate hydrogen bonds, while lines represent stacking interactions with DNA bases. The coloring corresponds to the major structural elements. **E.** A close-up view of the interactions between the C-tail of TFAM and mtRNAP in IC0. **F.** A close-up view of the TFB2M binding site on mtRNAP in IC0. Potential hydrogen bonds are indicated by blue dotted lines.

In contrast, Structure TWO revealed the GAG product bound to the T strand in the active site of mtRNAP, with the transcription bubble extending to 8 bp, spanning bases –4 to +4 (**Figure 1D**, **Figure 2B**). This complex, termed IC3, displayed a fully “open” conformation of the fingers domain, consistent with the post-translocated state in which the *n+1* templating base was unpaired and poised to bind the next incoming NTP substrate. The position of other important structural elements in mtRNAP, such as the intercalating hairpin and thumb subdomain, are nearly identical between IC0 and IC3 (**Figure 2 A,B**).

Unlike the previous pseudo-atomic model of the human mitochondrial initiation complex obtained through X-ray studies, the structures of IC0 and IC3 were resolved at near-atomic resolution (3.04 Å and 3.08 Å, respectively). These structures revealed a comprehensive network of interactions among the protein components of the initiation complex, the nucleic acid component, and the mechanism of promoter binding and recognition, as described below (**Figure 2 C,D, S3**).

### MtRNAP Interactions with TFAM and TFB2M in IC0 and IC3

IC0 and IC3 exhibit a more compact overall shape compared to previously reported structures (Hillen *et al*., 2017a), with all bases of the T and NT strands in the transcription bubble clearly resolved (**Figure 2 C, D, S3**). Despite their close proximity within the complex, TFAM and TFB2M do not interact directly with each other (**Figure 1 C,D**).

A key structural feature facilitating mtRNAP recruitment to the promoter is the tether helix in mtRNAP, which serves as the primary interface for TFAM binding (**Figure 2E**). The IC0 structure reveals close proximity of the “RKD” loop (residues 226–228), an important functional element of TFAM, to the loop-helix-loop motif (residues 145-162) in mtRNAP, which connects the PPR domain to the tether helix (**Figure 2E**). Specifically, in the IC0 structure, residue R226 of TFAM is in a position to interact with the carbonyl groups of residues 157 and 156 in mtRNAP. Residue D228 of the RKD loop is positioned to hydrogen bond with the hydroxyl group of T157, and a salt bridge is observed between E223 of TFAM and R159 of mtRNAP. These interactions, observed in the IC for the first time, explain the transcriptional deficiencies observed in TFAM variants lacking the RKD loop, which are incapable of supporting promoter-specific transcription (Dairaghi et al., 1995; Morozov *et al*., 2014; Shadel and Clayton, 1997).

The binding site for TFB2M is formed by two adjoining β-hairpins in mtRNAP: the intercalating hairpin (residues 605–624), which is directly involved in DNA melting, and the lever loop (residues 588–604), a structural feature unique to mitochondrial RNAPs and absent in phage polymerases (**Figure 2F**). Together, these loops form the binding interface with the C-terminus of TFB2M. However, this interaction is relatively weak, with only a few hydrophobic contacts and hydrogen bonds observed. Specifically, R601 and Y610 of mtRNAP form hydrogen bonds with the backbone carbonyl groups of P344 and the imidic nitrogen of R330 in TFB2M, respectively. The C-tail (res 385-396) of TFB2M inserts into a tight, ∼8 Å opening between the intercalating hairpin and the Thumb subdomain of mtRNAP (**Figure 2F**). Stabilization of the C-tail is achieved through a single hydrogen bond between T392 and R770 in the Thumb subdomain, along with additional interactions involving the NT strand of DNA, which are discussed below.

Overall, despite the extensive interaction interface between TFB2M and mtRNAP, most of the stabilizing interactions are mediated by contacts with the DNA component of the IC. This finding is consistent with the inability of TFB2M to bind mtRNAP directly in the absence of DNA (Morozov et al., 2015).

### Promoter Binding and Recognition

Recruitment of mtRNAP to promoter and formation of a pre-initiation complex is likely the first critical step in transcription initiation, allowing for positioning of the active site of polymerase over the TSS (Morozov *et al*., 2014; Morozov *et al*., 2015). In sharp contrast to the related single-subunit RNAP from bacteriophage T7, which recognizes bases in the major groove of the double-stranded promoter (Cheetham et al., 1999), mitochondrial promoter recognition by mtRNAP seems to rely on the melted region of DNA (**Figure 3**).

**Figure 3.**
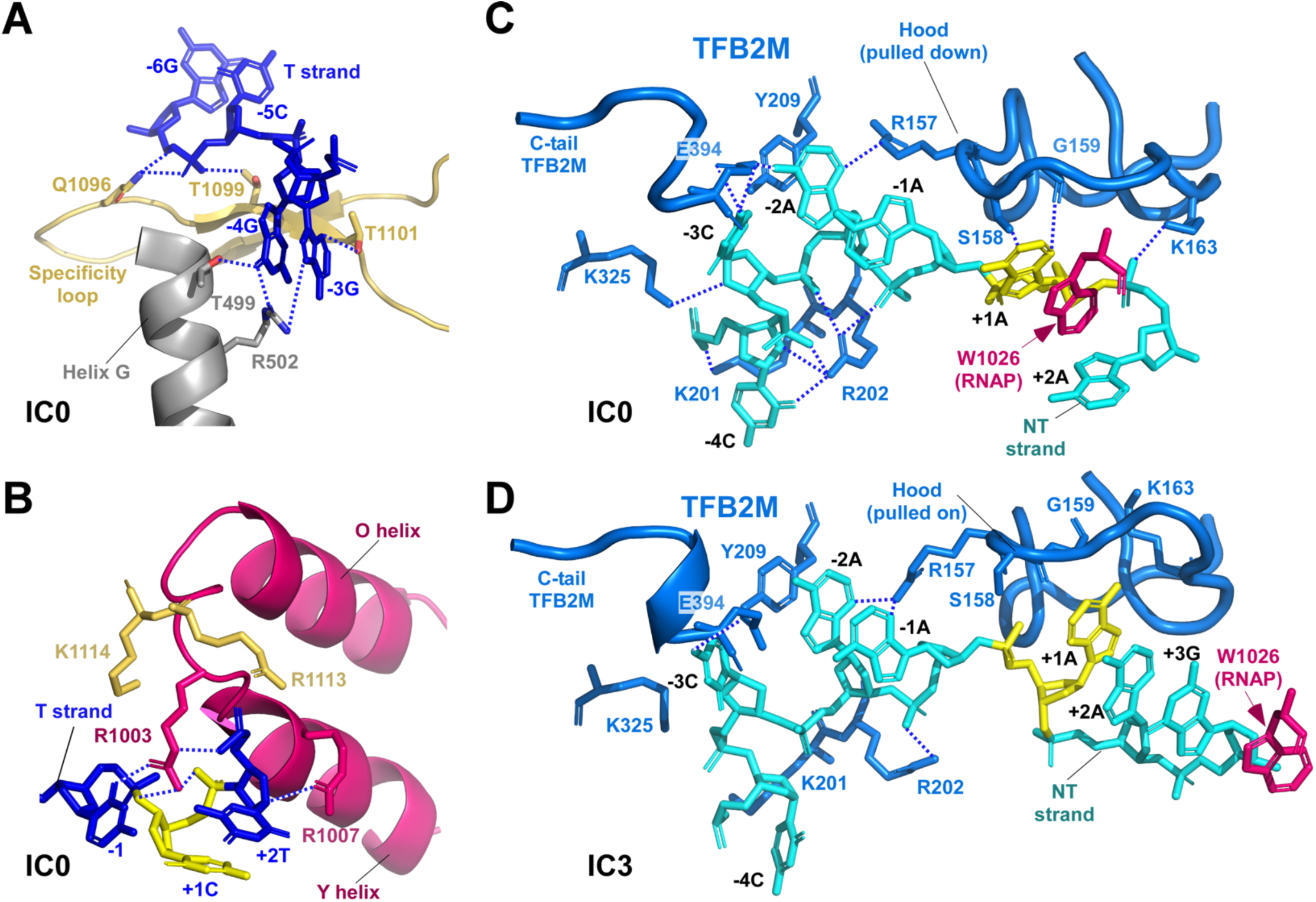
Mechanism of Promoter Recognition by the Human Transcription Initiation Complex. **A.** Recognition of the –4 and –3 bases in the T strand of the promoter in IC0. The –3 base (dAMP in the template used for the IC0/IC3 structures) is labeled as “-3G” due to the presence of dGMP at this position in human promoters. **B.** A close-up view of TSS in IC0. The O and Y helices of the fingers domain and residues in the base of the specificity loop (R1113 and K1114) are shown. The TSS base (+1) in the T strand is highlighted in yellow. Possible hydrogen bonds are indicated by blue dotted lines. **C,D.** Promoter recognition by TFB2M in IC0 (C) and IC3 (D). Possible hydrogen bonds are indicated by blue dotted lines. The TSS base (+1) in the NT strand is highlighted in yellow.

Three elements in mtRNAP provide a major contribution to promoter binding – the fingers domain, the intercalating hairpin, and the conserved in single-subunit RNAPs β-hairpin, referred to as the specificity loop (**Figure 3 A,B**).

Upon TFB2M binding, the intercalating hairpin is inserted between the DNA strands, juxtaposed to the –5 base pair, which remains within the DNA duplex. The intercalating hairpin establishes a number of interactions with the promoter, facilitating its binding and enabling the melting of the –4 and –3 bases (**Figure 2C,B**). These melted bases are subsequently recognized through interactions with helix G and the specificity loop (**Figure 3A**). Previous structural studies of human transcription initiation suggested that mtRNAP does not play a major role in promoter recognition (Hillen *et al*., 2017a). Indeed, the specificity loop is not fully inserted into the major groove of the promoter DNA and, therefore, does not engage in base-specific interactions with the –7 to –12 region of the promoter, as observed in the T7 RNAP IC (Cheetham and Steitz, 1999; Jeruzalmi and Steitz, 1998). Instead, it contributes to DNA binding by interacting with the phosphate backbone of DNA (**Figure 3A**).

Sequence-specific recognition of the conserved –4G and –3G bases within the LSP and HSP promoters is mediated by R502 of Helix G (**Figure 3A**). The side chain of R502 interacts with both guanine bases in the T strand, while recognition of O6 of –4G is aided by the hydroxyl group of T499. Additionally, the T1101 residue in the base of the specificity loop interacts with the –3G base in the T strand. These interactions are key to species-specific mitochondrial transcription, as demonstrated by comprehensive mutagenesis studies of human promoters (Zamudio-Ochoa *et al*., 2022). Furthermore, the substitution of these residues with the corresponding residues from porcine mtRNAP enables a switch in promoter specificity (Zamudio-Ochoa *et al*., 2022).

In IC0, the T strand is sharply kinked, forming an approximately 90° angle between the downstream DNA and the region containing the transcription bubble. This kink is stabilized by hydrogen bonding between the side chain of R1003 in the fingers domain of mtRNAP and the phosphate backbone at positions +1 and –1 of the T strand. The T strand path is additionally stabilized by interactions with the conserved residues, R1113 and K1114, located at the base of the specificity loop and carbonyl groups of R1003 and G1005 in the fingers domain (**Figure 3B**). The R1003 residue, which projects from the fingers domain towards the DNA and stacks upon the +1 base, likely plays a key role in enabling transcription initiation with purine nucleotides only (Zamudio-Ochoa *et al*., 2022), because the binding site for the +1 template base in IC0 can only accommodate a pyrimidine base. The conserved R1007 residue of the O-helix interacts with the +2T base in TS, explaining the conservation of this base in most mammalian species (**Figure 3B**).

The role of TFB2M in sequence-specific promoter recognition has been a long-standing question in the field (Gaspari et al., 2004; Matsunaga and Jaehning, 2004; Shadel and Clayton, 1995; Sologub *et al*., 2009). Recent studies indicated that TFB2M interacts with bases in the NT strand near the TSS, playing a critical role in species-specific promoter recognition (Zamudio-Ochoa *et al*., 2022). The structure of IC0 offers the first detailed insights into how TFB2M’s interactions enable promoter recognition (**Figure 3C, D**).

In IC0, TFB2M makes base-specific interactions with the NT strand using three distinct functional domains: the flexible loop in the N-terminal domain, referred to here as the “Hood”; the α5 helix (residues 200-213); and the C-terminal tail (residues 388–396) (**Figure 3C**). The Hood “covers” the +1 base in the NT strand, which is stabilized by stacking interactions with the conserved W1026 residue in mtRNAP, the only residue in mtRNAP that directly interacts with the NT strand of the promoter. Two residues in the Hood, S158 and G159, make hydrogen bonds with the +1 base via the hydroxyl group of S158 and the carbonyl group of G159. Interestingly, the Hood is absent in the yeast ortholog of TFB2M, Mtf1 (Goovaerts et al., 2023). Conversely, yeast mtRNAP lacks a residue in the fingers domain that corresponds to W1026 in the human polymerase, suggesting a different mechanism for transcription start site selection.

The –2A base in the NT strand is stabilized by stacking interactions with the Y209 residue and recognized by R157 (to N3 position) and the E394 (to N6) residues in the C-tail of TFB2M. The side chain of E384 residue also interacts with the N4 of the –3C base, which is stabilized by interaction with the carbonyl group of the same residue. Finally, the R202 residue of the a5 helix of TFB2M recognizes the O2 position of the –4C base in the NT strand (**Figure 3C**).

Both major human promoters are conserved at residues –4, –3, and +1; moreover, these positions in promoters are conserved in simians (Zamudio-Ochoa *et al*., 2022). Conversely, TFB2M residues identified in IC0 in direct interaction with these bases are also invariant in these species but differ in mammals of different orders (Zamudio-Ochoa *et al*., 2022).. Deletions or substitutions of the key residues in helix 5 and C-tail of TFB2M result in a significant loss of transcription stimulation by TFB2M (Hillen *et al*., 2017a).

### Initial RNA Synthesis

In IC3, the leading edge of the transcription bubble expands by 2 bp to accommodate the binding of the 3-mer RNA product (**Figure 2B,D**). The T strand of DNA descends into the active site, where it is positioned within the RNA-DNA binding cleft, while the NT strand undergoes scrunching. The interactions between mtRNAP and the –3G base are lost; however, a new network of interactions is formed between the bases near the TSS and the catalytic residues of mtRNAP (**Figure 2D**). The scrunching of the NT-strand, in turn, leads to a rearrangement of its interactions with TFB2M (**Figure 3D**).

Although several initial hydrogen bonds with TFB2M are lost during this process, NT DNA binding is stabilized by hydrophobic and van der Waals interactions. This stabilization involves the refolding of the Hood element, allowing the +1 base to insert into a newly formed hydrophobic cleft (**Figure 3D**). Due to the scrunching of the NT strand, the +1 base is no longer stacked with W1026, which instead interacts with the +3 base. The R202 residue disengages from the –4C base, while E394 remains hydrogen-bonded to the –3C base.

The continuous and tight interactions between TFB2M and the NT strand enable mtRNAP to maintain its position on the promoter and accommodate the 3-mer RNA product without translocating along the DNA or disrupting contacts with TFAM. This mechanism ensures the stability and fidelity of the transcription IC during *de novo* synthesis or the recycling of small RNA products.

### Transition to the Late-Stage Initiation, Promoter Clearance, and Initiation Factor Release

Since an artificially pre-melted promoter is known to interfere with promoter clearance (Temiakov et al., 2002), we employed fully double-stranded promoter templates for analysis of late-stage transcription initiation. Using a partial nucleotide mix, mtRNAP was allowed to synthesize an 8-mer RNA product in the presence of the GAG primer, ATP, and 3’dGTP (**Figure 4A**). Using SPA, we identified three conformations of the late IC complex, resulting in three structures that represent late stages of transcription initiation (**Figure 4B-D, S4-7, Table S2**).

**Figure 4.**
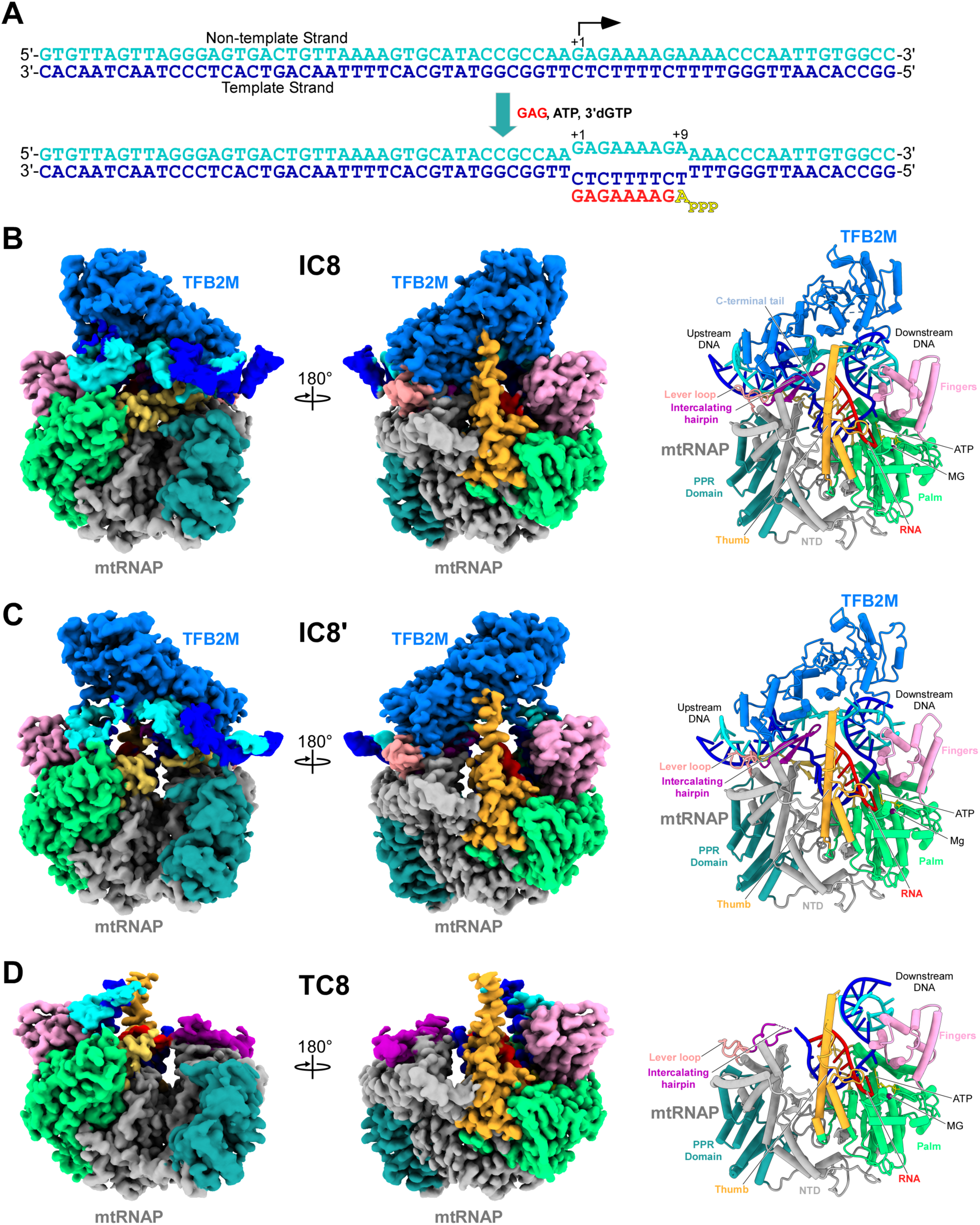
Structure of the Late-Stage Transcription Initiation Complex. **A.** Design of a fully double-stranded promoter template used for cryo-EM. The ICs were generated by transcription in the presence of GAG primer, ATP, and chain-terminating 3’dGTP. The incoming ATP observed in the insertion site of mtRNAP is indicated in yellow. **B.** The cryo-EM density and the structure of IC8. **C.** The cryo-EM density and the structure of IC8’. **D.** The cryo-EM density and the structure of the transitional complex, TC8.

In all the late-stage ICs, synthesis of the 8-mer RNA product has completed, resulting in a post-translocated state of mtRNAP and an incoming nucleotide bound in the Insertion Site of polymerase (**Figure 4 B-D**). In IC8 and IC8’, six nt of RNA are visible in the cryo-EM maps, while the density is weak for the two nt at the 5’ end of the chain (**Figure 5 A, B, S7**). Another structure, referred to as the Transitional Complex (TC8), represents an intermediate state between the IC and EC. The absence of upstream DNA density in the cryo-EM maps prevents its definitive classification as either an initiation or elongation complex. Notably, in TC8, all eight base pairs of the RNA-DNA heteroduplex are resolved in the cryo-EM maps (**Figure S7**).

**Figure 5.**
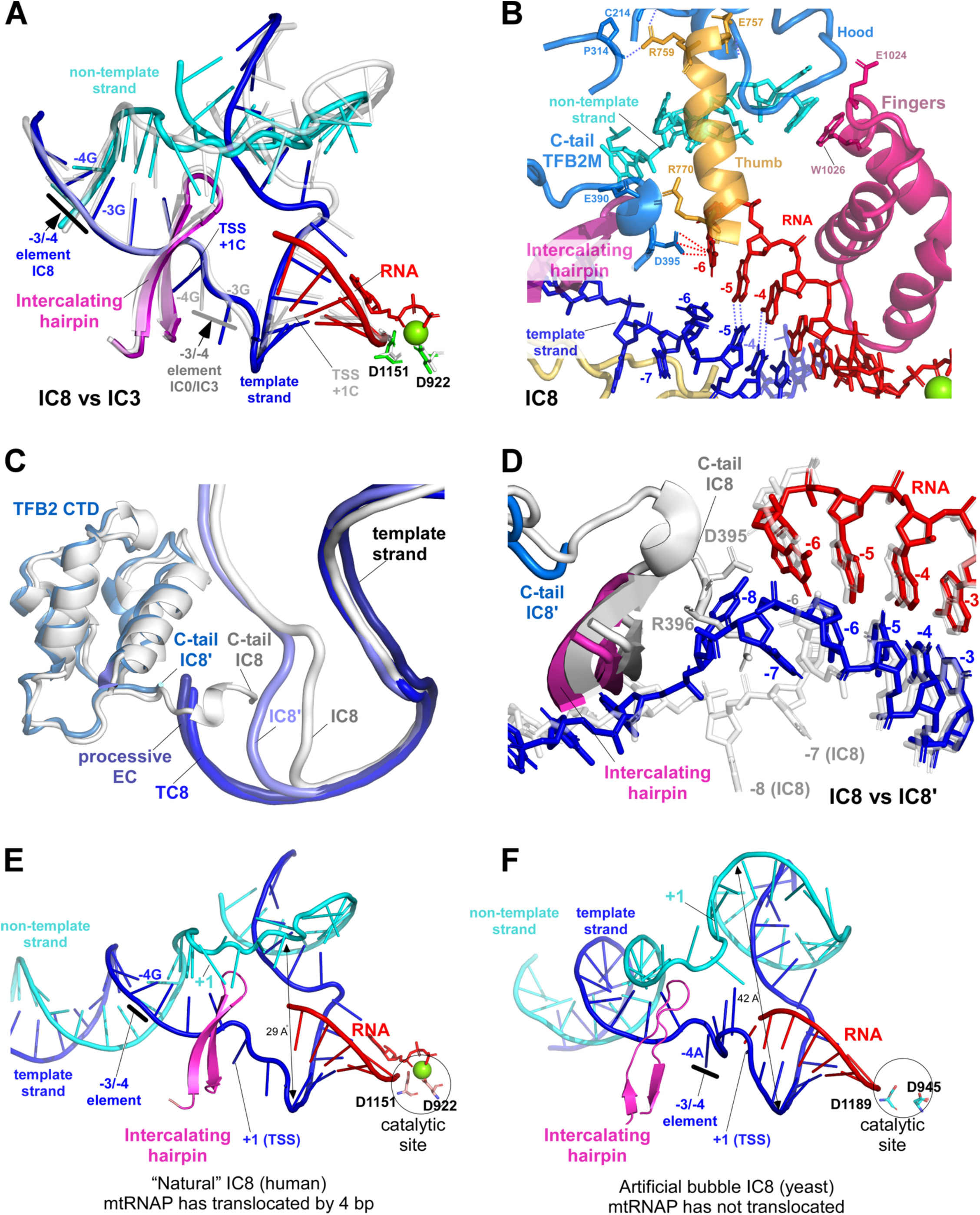
Late-Stage Transcription Initiation Complexes. **A.** Synthesis of 8-mer RNA requires only 4 translocation steps. IC3 and IC8 were aligned using residues in their conserved palm subdomain. The location of the –4/-3 recognition element in the promoter is indicated by a grey bar (IC3) and a black bar (IC8). **B.** A close-up view of the trailing edge of the transcription bubble in IC8. The proximity of residue D295 in the C-tail of TFB2M to the 5’ end of RNA is shown by dotted red lines. **C.** Changes in the trajectory of the T strand in the late-stage ICs. The trajectory of the T strand shifts during the transition from IC8 to IC8’ (light blue) and further in TC8 (dark blue). In TC8, the T strand trajectory closely matches that of the processive EC (PDB ID: 8U8V). **D.** The mechanism of TFB2M release. IC8 (white) and IC8’ were aligned using their corresponding palm subdomain. The C-tail of TFB2M in IC8 is observed clashing with the T strand in IC8’. **E,F.** Lack of DNA strand reannealing of the trailing edge of the artificial bubble in yeast IC8 complex. Nucleic acids in human IC8 (E) and yeast IC8 (F) are shown in the same orientation as their corresponding RNAPs. The location of the –4/-3 recognition element in the promoter is indicated by the black bar.

Superimposition of the IC8 with IC3 reveals that mtRNAP translocated four nucleotides along the DNA template, as evidenced by the positioning of the –4/-3 promoter element into the upstream region of the duplex DNA, “behind” the intercalating hairpin (**Figure 5A**). This finding suggests that the progression to IC8 begins after the incorporation of four nucleotides into the RNA transcript, corresponding to the IC4 stage. Each translocation step is accompanied by a ∼35° rotation of mtRNAP along the DNA duplex and a ∼6 Å movement of the tether helix away from the TFAM C-terminal domain, disrupting the hydrogen bonds that link these proteins. Consistent with this, the structures of the late-stage ICs reveal the absence of TFAM in cryo-EM maps and poorly resolved far-upstream DNA region (bases –17 to –40), indicating that mtRNAP-TFAM interactions are lost at this stage of transcription initiation. Given that even a single translocation step will disrupt the RKD loop interactions with mtRNAP, we speculate that TFAM dissociation occurs as early as the synthesis of a 4-5 mer RNA product (**Figure 5A**).

In IC8, the single-stranded region of the NT strand extends to nine base pairs, resulting in additional scrunching (**Figure 5A,B**). The flexibility of TFB2M enables it to accommodate all melted bases of the NT strand and make close interactions with the tip of the Thumb subdomain of mtRNAP, which becomes in the cryo-EM maps of IC8 (**Figure 5B**). The RNA-DNA hybrid in IC8 consists of five base pairs, while the –6 DNA base is flipped out, preventing Watson-Crick interaction with its RNA counterpart. The carboxyl group of residue D395 in the C-tail of TFB2M is positioned within 2.5 Å of the –6 RNA base, effectively blocking the further expansion of the RNA-DNA heteroduplex (**Figure 5B**).

In IC8, the trajectory of the T strand of DNA is nearly identical to its path in IC3 (**Figure 5C**). In contrast, in IC8’, the T strand adopts a conformation that more closely resembles the A-form of DNA, typical for RNA polymerases. Finally, in TC8, the trajectory of the T stand aligns closely with that of DNA in the processive EC (Hillen *et al*., 2017b; Schwinghammer et al., 2013) (**Figure 5C**), suggesting that these captured conformations of the late-stage ICs represent critical transitional steps.

The altered trajectory of the T strand in IC8’, along with an expansion of the RNA-DNA heteroduplex from five to six base pairs, creates a steric clash with the C-tail of TFB2M (**Figure 5D**), suggesting a mechanism for its displacement. Superimposition of IC8 and IC8’ indicates that in IC8’, the position of the C-tail is now occupied by the –8 and –7 bases of DNA, resulting in the disappearance of the terminal eight residues of TFB2M from the cryo-EM maps (**Figure 5D**). Further expansion of the RNA-DNA heteroduplex in TC8 and steric clashes with TFB2M likely drive its complete displacement from the IC, facilitating the transition to the elongation stage of transcription (**Figure 5 C, D**).

The structures of late-stage human mitochondrial ICs formed on fully double-stranded promoters reveal a mechanism by which mtRNAP maintains the size of the transcription bubble. This mechanism involves a “zipping” action mediated by the intercalating hairpin and relies on the complementary NT strand as a driving force for DNA strand reannealing at the trailing edge of the transcription bubble. In the absence of a complementary NT strand in the upstream region, mtRNAP fails to translocate during the synthesis of the 8-nt RNA product (Goovaerts *et al*., 2023). This failure leads to severe scrunching of both strands of the promoter DNA and an inability of the IC to clear the promoter and transition to the elongation phase of transcription, as observed recently in yeast mitochondrial transcription ICs formed on pre-melted promoter DNA (**Figure 5 E,F**) (Goovaerts *et al*., 2023).

### Transition to the Elongation Stage of Transcription

To probe at which stage of transcription initiation mtRNAP transitions into a processive EC, we initiated transcription on a fully double-stranded promoter template in the presence of the elongation factor, TEFM (**Figure S8**). The binding of TEFM is incompatible with the topology of the DNA in the early-stage IC and requires TFB2M dissociation (Hillen *et al*., 2017b). In the presence of GAG primer, ATP, and 3’dGTP, mtRNAP was allowed to synthesize a 9-mer RNA product (**Figure 6A, S8 A,B, S9**). In addition, to probe the structure of EC formed in the absence of TEFM, we generated a promoter-originated EC13 (**Figure 6B, S8 C,D, S10**).

**Figure 6.**
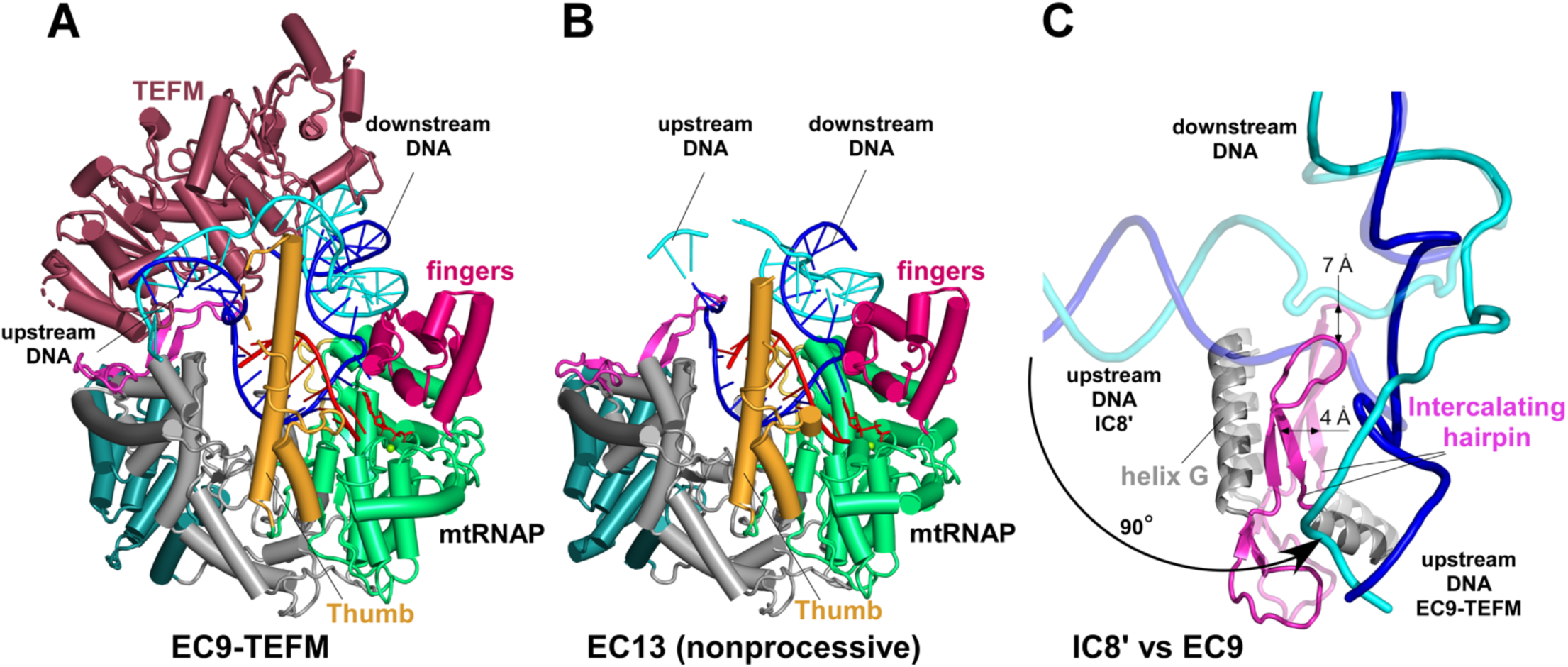
Transition from Initiation to Elongation. **A.** Structure of the promoter-originated EC9-TEFM complex. The complex was generated in the presence of TFAM, TFB2M, and the elongation factor, TEFM. **B.** Structure of a nonprocessive promoter-originated EC13. **C.** Changes in the trajectory of the upstream DNA during the transition from IC to EC. IC8’ and EC9 structures were superimposed using their palm subdomains.

SPA of promoter-originated EC9 revealed particles lacking TFB2M but containing TEFM, indicating the formation of a processive EC-TEFM complex (**Figure 6A, Figure S9, S11**). The overall topology of the nucleic acid components and proteins in EC9-TEFM and EC13 is nearly identical to the previously determined structure of ECs assembled on nucleic acid scaffolds (Cα root-mean-square deviation [RMSD] <1 Å over 560 residues as compared to PDB 8U8V) (**Figure 6A, B, S12**) (Herbine et al., 2024; Hillen *et al*., 2017b).

Superimposition of EC9-TEFM onto IC8’ reveals a ∼90° shift in the axis of the downstream DNA duplex (**Figure 6C**), accompanied by relative movements of the N-terminal domain and C-terminal domain of mtRNAP. This movement “opens” the heteroduplex binding cavity of mtRNAP and induces a marked shift in the intercalating hairpin (**Figure 6C**), which translates ∼4 Å away from the CTD and ∼7 Å away from the upstream DNA duplex. This shift allows the upstream DNA duplex in IC8’ to “swing” over the tip of the intercalating hairpin and adopt a more relaxed conformation observed in all ECs. The shift in the upstream DNA trajectory during the transition from IC to EC is independent of TEFM, as similar conformational changes in mtRNAP, including intercalating hairpin movement, are observed in EC13 formed without TEFM (**Figure 6**).

## DISCUSSION

### The sequence of transcription events and a comprehensive model of transcription initiation

The structural analysis of early– and late-stage transcription initiation complexes provides a detailed reconstruction of the sequence of events in mitochondrial transcription (**Figure 7, Video S1**). Recruitment of mtRNAP to the promoter through interactions with TFAM induces conformational changes in the TFB2M binding site (Hillen *et al*., 2017a), enabling TFB2M to load onto the pre-initiation complex. Once bound, TFB2M facilitates the initial melting of six base pairs of promoter DNA, as observed in IC0, and establishes interactions with both strands of the melted DNA region through TFB2M and mtRNAP (**Figure 3C,D)**.

**Figure 7.**
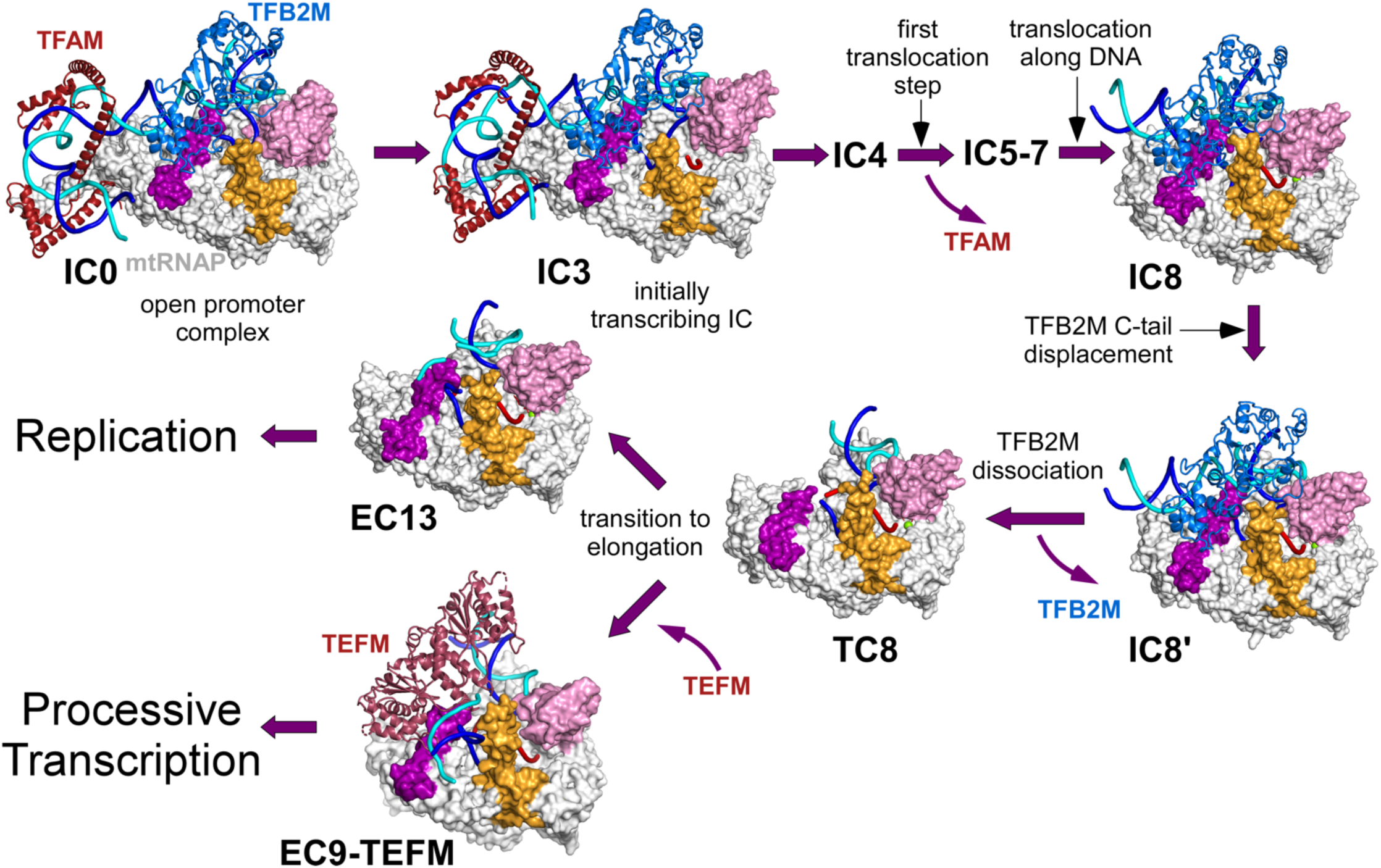
Structural basis for transcription initiation, promoter clearance, and transition to elongation. Major steps of transcription initiation at the LSP promoter are indicated. Structures are shown in the same oientation as heir conserved palm subdomains in mtRNAP.

The structure of IC0, a catalytically inactive complex, provides critical insights into how mammalian promoters are recognized through the coordinated action of its three protein components. While extensive biochemical and genetic studies over the years have advanced our understanding of these processes, the precise mechanistic roles of TFAM, mtRNAP, and TFB2M remained unclear (Gaspari *et al*., 2004; Hillen *et al*., 2017a; Morozov *et al*., 2014; Morozov *et al*., 2015; Morozov and Temiakov, 2016; Ngo et al., 2014; Rubio-Cosials and Sola, 2013; Zamudio-Ochoa *et al*., 2022). The structural information from IC0 offers a detailed view of the interactions occurring during the initial stage of initiation, revealing for the first time the roles of TFB2M, and the specificity loop and fingers domain of mtRNAP in DNA binding and recognition. These findings explain the phenotypes of numerous previously characterized variants of mtRNAP, TFAM, and TFB2M.

The binding of substrate NTPs or short RNA primers facilitates the transition of IC0 into the catalytically competent IC3. Structural comparisons between IC0 and IC3 indicate that this step occurs without mtRNAP translocation, as the polymerase maintains its interactions with the upstream promoter and TFAM. Since the templating DNA base is already melted in IC3, the transition to IC4 and the synthesis of a 4-mer RNA primer likely proceed without translocation, preserving TFAM association (**Figure 2B**). In contrast, the subsequent steps of transcription initiation, as observed in IC8 structures, involve mtRNAP translocating along the DNA template (**Figure 5A**). This movement disrupts the interactions between TFAM and mtRNAP, signaling the transition toward elongation.

The IC structures reveal a fundamental difference between the transition mechanisms in mitochondrial RNAPs and T7 RNAP, a widely used model for single-subunit RNAPs. In T7 RNAP, the synthesis of RNA transcripts longer than 3 nucleotides triggers a dramatic refolding of the N-terminal domain, which otherwise obstructs the RNA-DNA hybrid’s path in the heteroduplex binding cavity (Tahirov et al., 2002; Yin and Steitz, 2002). This refolding process is believed to cause the synthesis of numerous abortive RNA products (Ramirez-Tapia and Martin, 2012; Vahia and Martin, 2011). In contrast, human mtRNAP follows a distinct mechanism. The path of the T strand of DNA remains nearly identical in IC3 and IC8, with no evidence of T strand scrunching or major structural rearrangements in the polymerase (**Figure 5A**). This suggests minimal destabilizing forces during these stages of RNA synthesis. As a result, mtRNAP produces significantly fewer abortive products compared to T7 RNAP (Lodeiro et al., 2012; Sologub *et al*., 2009).

The key transitional step in mtRNAP initiation occurs when the RNA reaches a length of 8–9 nucleotides (**Figure 7**). At this stage, at the trailing end of the RNA-DNA hybrid, the T-strand forms a 5-6 bp heteroduplex that clashes with the C-tail of TFB2M, displacing it from the opening between the intercalating hairpin and the Thumb subdomain, as observed in IC8’ structure (**Figure 5D**). This displacement leads to the dissociation of TFB2M (**Figure 7**).

The loss of TFB2M disrupts mtRNAP’s ability to maintain the strained conformation of the promoter DNA, where the upstream and downstream DNA axes are nearly parallel (**Figure 5A, S13**). Conformational changes in mtRNAP follow, marked by a relative movement of its N-terminal and C-terminal domains. These changes result in the removal of the intercalating hairpin from its position between the DNA strands in IC8/IC8’, allowing the upstream DNA duplex to swing ∼90° into its final position within EC (**Figure 6C**). In this conformation, the intercalating hairpin now controls the size of the RNA-DNA hybrid. These structural rearrangements complete the isomerization of the IC into an EC (**Figure 7**).

Our data suggest that TEFM recruitment can occur as early as when mtRNAP begins synthesizing an 8-9 nt RNA. At this stage, TFB2M has dissociated from mtRNAP, freeing the binding site for TEFM, as observed in TC8 structure (**Figure 4D**). We propose that TEFM binding to the EC is stochastic, resulting in two potential outcomes: the ECs with TEFM become processive, while others, having no bound TEFM, remain nonprocessive, either dissociating or producing short RNA primers required for replication (**Figure 7**). The stochastic mode of TEFM binding likely represents the initial step in the model of the transcription-replication switch, which serves to prevent harmful collisions between replication and transcription machinery (Agaronyan *et al*., 2015; Gupta et al., 2023). Additional regulatory steps, such as replisome reloading at the end of the D-loop region, are likely required to fully execute and control this switch (Jemt et al., 2015).

In conclusion, this study provides a comprehensive structural and mechanistic framework for understanding transcription initiation in human mitochondria, highlighting the intricate coordination between mtRNAP and transcription factors. By resolving early– and late-stage initiation complexes, we have reconstructed the sequential events of transcription initiation, from promoter recognition and melting to RNA synthesis and the transition into elongation. The findings illuminate the unique strategies employed by mtRNAP, distinct from other single-subunit RNAPs, to minimize abortive RNA synthesis and maintain transcriptional fidelity.

## Acknowledgments

We thank current and former members of the Temiakov laboratory. We are indebted to Mahira Aragon, Alex (Hui) Wei, Aygul Ishemgulova, Jing Wang, and Kasahun Neselu (NCCAT) for their expert technical assistance during data collection. We also thank Dr Gino Cingolani for his expert suggestions and fruitful discussion.

## Funding

National Institutes of Health grant R35 GM131832 (DT). Some of this work was performed at the National Center for CryoEM Access and Training (NCCAT) and the Simons Electron Microscopy Center located at the New York Structural Biology Center, supported by NIH (Common Fund U24GM129539, NIGMS R24GM154192), the Simons Foundation (SF349247) and NY State Assembly. Initial cryo-EM analysis was carried out at Jefferson Integrate Structural Biology Shared Resources, which is supported in part by grant S10 OD030457.

## Author contributions

Experimental design and conceptualization: KH, ARN, DT. Protein preparation and biochemical experiments: KH Cryo-EM Data acquisition: KH Single-particle Cryo-EM analysis and model building: KH, ARN Protein and DNA alignment and analysis: AZO

Writing – original draft: DT Writing – review & editing: DT, KH, ARN, AZO

## Competing interests

The authors declare no competing interests.

## Data and materials availability

The Cryo-EM maps and atomic coordinates were deposited in the Electron Microscopy Data Bank under accession codes EMD-48413, EMD-48412, EMD-48414, EMD-48415, EMD-48416, EMD-48417, and EMD-

48417, and in the Protein Data Bank under accession codes 9MN5, 9MN4, 9MN6, 9MN7, 9MN8, 9MN9, and 9MNA.

## Supplemental Information

Supplemental Information includes figures S1-S13, tables S1-S3, and Video S1.

## Material and Methods

### Expression of the components of human mitochondrial transcription components

A variant of human mtRNAP lacking the N-terminal 119 amino acids (Δ119 mtRNAP) with an N-terminal 6xHIS tag was constructed from the pProEx-based expression plasmid and described previously (Morozov *et al*., 2014). N-his TFAM lacking the mitochondrial localization sequence (Δ42 TFAM) and two cysteine residues (C49S and C246S) was expressed using the pET22b-based plasmid as described previously (Morozov *et al*., 2014). Human N-his TFB2M was expressed using a construct lacking a putative mitochondrial localization sequence (Δ20 TFB2M) (Sologub *et al*., 2009). Proteins were expressed in *E.coli* BL21 (DE3) RIL cells (Agilent).

### Purification of Mitochondrial RNA Polymerase

To express mtRNAP, the cells were grown in LB media containing 50 µg/ml Ampicillin, 50 µg/ml Chloramphenicol, and 0.8% D-glucose at 37 °C and shaken at 160 rpm until the OD_600_ reached 0.6 units. Expression was induced by the addition of 0.15 mM IPTG and carried out at 16 °C for 18 h. The cells were harvested by centrifugation at 8,000 g for 20 min and resuspended in mtRNAP lysis buffer. The cells were lysed by ultrasonication (pulsed for 1s on and 2s off) for 20 minutes and the cell debris was pelleted by centrifugation at 18,000 g for 30 min, twice. Soluble 6xHis-tagged mtRNAP, was loaded on to Ni-NTA beads (QIAGEN) pre-equilibrated with mtRNAP Ni-NTA wash buffer in a PD-10 Sephadex G-25 resin column (Cytiva Life Sciences) and eluted with mtRNAP Ni-NTA elution buffer. The protein was further purified by Heparin affinity chromatography using a HiTrap Heparin HP column (GE Healthcare). The heparin column was equilibrated in mtRNAP Heparin Buffer A and was eluted by a 0-70% linear gradient of mtRNAP Heparin Buffer B in a total of 8 column volumes. Peak fractions were collected and analyzed using SDS-PAGE. If necessary, mtRNAP peak fractions were pooled and dialyzed overnight to prepare for further purification by size exclusion chromatography which was performed using a Superdex 200 Increase 10/300 column (GE Healthcare) equilibrated with mtRNAP size exclusion buffer. The final combined fractions were applied to a 4 mL centrifuge concentrator with a membrane cut off of 50 kDa (Amicon) to 8-11 mg/mL at 4,000 g at 16 °C. The final protein preparation was aliquoted and stored at –80 °C.

### Purification of TFAM

The cells were grown in LB media containing 50 µg/ml Ampicillin, 50 µg/ml Chloramphenicol, and 0.8% D-glucose at 37 °C and shaken at 160 rpm until the OD_600_ reached 1.0 unit. Expression was induced by addition of 0.4 mM IPTG and carried out at 37 °C for 2 h. The cells were harvested by centrifugation at 8,000 g for 20 min and resuspended in TFAM lysis buffer. The cells were lysed by ultrasonication (pulsed for 30s on and 40s off) for 20 minutes and the cell debris was pelleted by centrifugation at 27,000 g for 30 min, twice. Soluble 6xHis-tagged TFAM was loaded on to Ni-NTA beads (QIAGEN) pre-equilibrated with TFAM Ni-NTA wash buffer in a PD-10 Sephadex G-25 resin column (Cytiva Life Sciences) and eluted with TFAM Ni-NTA elution buffer.

TFAM was further purified by Heparin affinity chromatography using a HiTrap Heparin HP column (GE Healthcare). The heparin column was equilibrated in TFAM Heparin buffer A and was eluted by a 0-80% linear gradient of TFAM Heparin Buffer B in a total of 10 column volumes. Peak fractions were collected and analyzed using SDS-PAGE. The selected fractions were dialyzed overnight at 4 °C against 1 L of TFAM Dialysis Buffer to prepare for the final step of purification. Dialyzed TFAM was purified using cation exchange chromatography on a MonoS column (GE Healthcare). The MonoS column was equilibrated in TFAM MonoS Buffer A and was eluted by a 0-75% linear gradient of TFAM MonoS Buffer B in a total of 15 column volumes. Peak fractions were analyzed by SDS-PAGE. The final combined fractions were applied to a 4 mL centrifuge concentrator with a membrane cut off of 10 kDa (Amicon) to 9-11 mg/mL at 4,000 g at 16 °C. The final protein preparation was aliquoted and stored at –80 °C.

### Purification of TFB2M

The cells were grown in LB media containing 50 µg/ml Ampicillin and Chloramphenicol at 37 °C and 160 rpm until the OD_600_ reached 3.0 units. Expression was induced by addition of 0.15 mM IPTG and carried out at 16 °C for 18 h. The cells were harvested by centrifugation at 8,000 g for 20 min and resuspended in TFB2M lysis buffer. The cells were lysed by ultrasonication (pulsed for 0.4 s on and 1 s off) for 20 minutes and the cell debris was pelleted by centrifugation at 18,000 g for 30 min, twice. Soluble 6xHis-tagged TFB2M was loaded on to Ni-NTA beads (QIAGEN) pre-equilibrated with TFB2M Ni-NTA wash buffer in a PD-10 Sephadex G-25 resin column (Cytiva Life Sciences) and eluted with TFB2M Ni-NTA elution buffer. TFB2M protein was further purified by Heparin affinity chromatography using a HiTrap Heparin HP column (GE Healthcare). The heparin column was equilibrated in TFB2M Buffer A and TFB2M was eluted by a 0-80% linear gradient of TFB2M Buffer B in a total of 10 column volumes. Peak fractions were collected and analyzed using SDS-PAGE. If necessary, TFB2M peak fractions were pooled and dialyzed overnight to prepare for further purification by size exclusion chromatography which was performed using a Superdex 200 Increase 10/300 column (GE Healthcare) equilibrated with TFB2M size exclusion buffer. The final combined fractions were applied to a 4 mL centrifuge concentrator with a membrane cut off of 10 kDa (Amicon) to 9-11 mg/mL at 4,000 g at 16 °C. The final protein preparation was aliquoted and stored at –80 °C.

### Purification of TEFM

N-terminal His-tagged Human TEFM (Δ50) was expressed in *E.coli* Rosetta2 (DE3) cells (Merck Millipore) and purified as described previously (Herbine *et al*., 2024). The cells were grown in LB media containing 50 µg/ml Ampicillin, 50 µg/ml Chloramphenicol, and 0.8% D-glucose at 37 °C and 160 rpm until the OD_600_ reached 0.6 units. Expression was induced by addition of 0.15 mM IPTG and carried out at 16 °C for 18 h. The cells were harvested by centrifugation at 8,000 g for 20 min and resuspended in TEFM lysis buffer. The cells were lysed by ultrasonication (pulsed for 0.4 s on and 1 s off) for 15 minutes and the cell debris was pelleted by centrifugation at 18,000 g for 30 min, twice. Soluble 6xHis-tagged TEFM was loaded on to Ni-NTA beads (QIAGEN) pre-equilibrated with TEFM Ni-NTA wash buffer in a PD-10 Sephadex G-25 resin column (Cytiva Life Sciences) and eluted with TEFM Ni-NTA elution buffer. TEFM was further purified by Heparin affinity chromatography using a HiTrap Heparin HP column (GE Healthcare). The heparin column was equilibrated in TEFM Heparin buffer A and was eluted by a 0-70% linear gradient of TEFM Heparin Buffer B in a total of 10 column volumes. Peak fractions were collected and analyzed using SDS-PAGE. The selected fractions were dialyzed overnight at 4 °C against 1 L of TEFM Dialysis Buffer to prepare for the final step of purification. Dialyzed TEFM was purified using cation exchange chromatography on a MonoS column (GE Healthcare). The MonoS column was equilibrated in TEFM MonoS Buffer A and was eluted by a 0-75% linear gradient of TEFM MonoS Buffer B in a total of 15 column volumes. Peak fractions were analyzed by SDS-PAGE. The final combined fractions were applied to a 4 mL centrifuge concentrator with a membrane cut off of 10 kDa (Amicon) to 9-11 mg/mL at 4,000 g at 16 °C. The final protein preparation was aliquoted and stored at –80 °C.

### DNA and RNA oligonucleotides and template annealing

All DNA oligos were from IDT DNA. The sequence of the oligos is below (5’ to 3’):

NT-50bub-3/+1: GAAAATAATGTGTTAGTTGGGGGGTGACTGTTAAAAGTGCATACCGCCAAAAGATAGGCC

TS-50bub-3/+1 GGCCTATCTCCCAGCGGTATGCACTTTTAACAGTCACCCCCCAACTAACACATTATTTTC

NT66_IC18: GTGTTAGTTAGGGAGTGACTGTTAAAAGTGCATACCGCCAAGAGAAAAGAAAACCCAATTGTGGCC

TS66_IC18: GGCCACAATTGGGTTTTCTTTTCTCTTGGCGGTATGCACTTTTAACAGTCACTCCCTAACTAACAC

NT66_IC10: GTGTTAGTTAGGGAGTGACTGTTAAAAGTGCATACCGCCAAGAGAAAAAGAAAACCCAATTGTGG CC

TC66_IC10: GGCCACAATTGGGTTTTCTTTTTCTCTTGGCGGTATGCACTTTTAACAGTCACTCCCTAACTAACAC

The RNA oligonucleotide, GAG, was from Dharmacon.

### Assembly of human mitochondrial ICs on pre-melted promoter DNA

To assemble the early-stage human mitochondrial initiation complexes, 20 µM Δ119 mtRNAP was mixed with Δ42 TFAM, Δ20 TFB2M, and a promoter bubble template (cryo_wt_NT-50bub LSP/cryoTS-50bub-3/+1 LSP/GAG) at a 1:3:2:1 molar ratio in the dialysis buffer containing 20 mM Tris-HCl (pH 7.9), 150 mM NaCl, 10 mM DTT, and 10 mM MgCl_2_ and put on overnight dialysis at 4 °C.

### Assembly of human mitochondrial ICs on fully complementary promoter templates

To generate late-stage human mitochondrial initiation complexes (IC8), 5 µM Δ119 mtRNAP was mixed with Δ42 TFAM, Δ20 TFB2M, and a promoter template (NT66_IC18/ TS66_IC18) at a 1:3:2:1 molar ratio in the dialysis buffer. Following the overnight dialysis, transcription was initiated by the addition of the GAG RNA primer (50 μM), ATP (125 μM), and 3’-deoxy GTP (250 μM) for 20 min at 25 °C.

To generate promoter-initiated human mitochondrial elongation complexes with TEFM (EC9-TEFM), 5 µM Δ119 mtRNAP was mixed with Δ42 TFAM, Δ20 TFB2M, Δ50 TEFM and a promoter template (NT66_IC10/ TS66_IC10) at a 1:3:2:2:2 molar ratio in the dialysis buffer. Following overnight dialysis, transcription was initiated by the addition of the GAG RNA primer (50 μM), ATP (125 μM), and 3’-deoxy GTP (250 μM) for 20 min at 25 °C.

To generate EC13, transcription was performed as described for EC9 above but in the absence of TEFM. RNA synthesis was initiated by the addition of the GAG RNA primer (50 μM), ATP (125 μM), GTP (125 μM), and 3’-deoxy CTP (125 μM) for 20 min at 25 °C.

### Cryo-EM grid preparation for early-stage transcription ICs

Cryo-EM grids were prepared at the National Center for Cryo-EM Access and Training (NCCAT) using a Chameleon (SPT Labtech) blot-free vitrification system. Self-wicking nanowire grids, Carbon 300 mesh grids (SPT Labtech) were negatively glow-discharged with 15 mA for 80 seconds using a PELCO easiGlow™ Glow Discharge Cleaning System prior to Chameleon use. 15 nl of sample was sprayed onto the grids with plunge times between 156 and 161 milliseconds before immediate freezing in liquid ethane. The sample quality and particle distribution were assessed using Leginon and Appion web tools (NCCAT) operating a 200kV Glacios™ cryo-TEM (ThermoFisher Scientific) equipped with a Falcon 4 direct electron detector.

### Cryo-EM grid preparation for late-stage ICs and ECs

Cryo-EM grids were prepared at Jefferson Cryo-EM facility. For preparations involving IC8, EC9-TEFM, and EC13 UltrAuFoil® R1.2/1.3, Gold 300 mesh grids (Quantifoil, Germany) were negatively glow-discharged with 15 mA for 90 seconds using a PELCO easiGlow™ Glow Discharge Cleaning System prior to preparation. The sample (3 µL) was then applied to grids that were blotted with PELCO® qualitative cellulose filter paper (Grade 595 55/20mm) and vitrified with a Vitrobot Mark IV (ThermoFisher Scientific, USA) for 5 seconds at 4°C and 95-100% humidity. The sample quality and particle distribution were assessed using EPU (Thermo Fisher Scientific) operating a 200kV Glacios™ cryo-TEM (ThermoFisher Scientific) equipped with a Falcon 4 direct electron detector.

### Cryo-EM data acquisition and image processing for ICs and ECs

Data was collected at the National Center for Cryo-Electron Microscopy Access & Training (NCCAT) at New York Structural Biology Center (NYSBC) using a 300 kV Titan Krios™ transmission electron microscope (ThermoFisher Scientific), equipped with a Falcon 4i direct electron detector and a Selectris Energy Filter using a 10 eV slit width. For early-stage ICs, movies were recorded in counting mode using Leginon^5^ (Suloway et al., 2005) with a magnification of 165,000, corresponding to a pixel size of 0.7511 Å for (micrograph dimensions of 4,096 × 4,096 px) over a defocus range of –0.3 to –2.5 µm. A dose rate of 7.95 e^-^/ Å^2^/s resulted in a total electron dose of 47.69 e^-^/Å^2^, which was averaged over 60 frames. A total of 26,965 exposures were collected and saved in electron-event representation (EER) format. For IC8, movies were recorded in counting mode using Leginon with a magnification of 165,000, corresponding to a pixel size of 0.7304 Å for (micrograph dimensions of 4,096 × 4,096 px) over a defocus range of –0.4 to –3.2 µm. A dose rate of 7.57 e^-^ /Å^2^/s resulted in a total electron dose of 45.44 e^-^/Å^2^, which was averaged over 60 frames. A total of 17,559 exposures were collected and saved in EER format. For EC9-TEFM, movies were recorded in counting mode using Leginon with a magnification of 165,000, corresponding to a pixel size of 0.7304 Å for (micrograph dimensions of 4,096 × 4,096 px) over a defocus range of –0.4 to –2.7 µm. A dose rate of 7.75 e^-^ /Å^2^/s resulted in a total electron dose of 46.52 e^-^/Å^2^, which was averaged over 60 frames. A total of 12,506 exposures were collected and saved in EER format.

For EC13, movies were recorded in counting mode using Leginon with a magnification of 165,000, corresponding to a pixel size of 0.7304 Å (micrograph dimensions of 4,096 × 4,096 px) over a defocus range of –0.3 to –2.2 µm. A dose rate of 7.42 e^-^ /Å^2^/sec resulted in a total electron dose of 45.44 e^-^/Å^2^, which was averaged over 60 frames. A total of 12,245 exposures were collected and saved in EER format. All movies were imported and processed in CryoSPARC (Punjani et al., 2017).

### Single Particle Analysis of early-stage human mitochondrial transcription IC

The workflow for image processing is shown in **Figure S1**. The EER exposures were imported and processed in CryoSPARC (Punjani *et al*., 2017) with EER fractionation set to 50 and the EER upsampling factor set to 1. The movies were converted into image stacks and were motion-corrected, gain-normalized, drift-corrected, summed, and dose-weighted using the Patch Motion Correction module (CryoSPARC). The Contrast transfer function (CTF) values of the motion corrected movies were estimated using Patch CTF (CryoSPARC). Micrographs with ice, ethane contamination, poor CTF fit resolution (>4 Å), and high astigmatism (≥ 1000 Å) were discarded. Particles were picked and extracted from the dose-weighted images with box size of 384 px (Fourier-cropped to 128 px) using CryoSPARC blob picker with dimensions of 80-160 Å and local maxima of 1000 picks. The entire dataset consisted of 24,411 motion-corrected images with 7,863,657 initial particles. Particles were aligned and sorted using two rounds of CryoSPARC 2D classification (n = 50), resulting in 4,957,265 curated particles. Initial models were generated using CryoSPARC *ab initio* reconstruction (n = 3) using a subset of the particles (1,000,000 particles) to generate initial reference-free maps. The *ab initio* reconstruction generated two maps representing the IC (1,2) and one “junk” particle class (3) that were used to curate all extracted particles within the dataset (4,957,265 particles) using heterogeneous refinement (n=6) using maps 1 and 2 as references and four “junk” classes used as decoys for particle sorting **(Figure S1)**. After three rounds of heterogeneous refinement,

2,441,562 particles representing the IC were refined using CryoSPARC non-uniform 3D refinement (Punjani et al., 2020) (128 box size) and aligned for downstream processing. To assess the conformational landscape of the data, CryoSPARC 3D Variability Analysis (3DVA) (Punjani and Fleet, 2021) was performed (n=5 modes, filter resolution limit 6 Å). The variability components, 0 and 1, estimated from the consensus non-uniform 3D refined map was dominated by motions in the fingers domain of mtRNAP and the upstream DNA region bound by TFAM, respectively. These components were passed through 3D Variability Display (Punjani and Fleet, 2021)(Simple mode, n=20 frames) and the maps were assessed for quality to generate 6 input maps (three maps from each component) for 3D classification. To probe the 3D variability results, we performed 3D classification (input mode, n=6, 6 Å filter resolution) on the particles to generate 6 distinct classes. Particles within classes 2-5 were initially explored with downstream heterogeneous and non-uniform 3D refinement but were ultimately rejected due to preferential orientation exhibited by the particles and poor resolution in regions of interest. Particles in class 1 represent IC0, contained 380,094 particles and was further processed by two rounds of heterogeneous refinement (n=2, force hard classification) using class 1 and a decoy class (class 3) as input volumes. A final non-uniform refinement (n=2 extra final passes, minimize per-particle scale) of class 1 resulted in Structure One with the following particle count and nominal resolution: IC0 (131,267 particles; 3.04 Å). Particles in class 6 represent IC3, contained 506,855 particles and was further processed by two rounds of heterogeneous refinement (n=2, force hard classification) using class 6 and a decoy class (class 3) as input volumes. A final non-uniform refinement (n=2 extra final passes, minimize per-particle scale) of class 6 resulted in Structure Two with the following particle count and nominal resolution: IC3 (109,470 particles; 3.08 Å).

### Single Particle Analysis of late-stage human mitochondrial transcription IC

Workflows for image processing of Complex TWO are found in **Figure S4**. The EER exposures were imported and processed in CryoSPARC with EER fractionation set to 60 and the EER upsampling factor set to 1. The movies were converted into image stacks and were motion corrected, gain-normalized, drift-corrected, summed, and dose-weighted using the Patch Motion Correction module (CryoSPARC). The Contrast transfer function (CTF) values of the motion corrected movies were estimated using Patch CTF (CryoSPARC). Micrographs with ice, ethane contamination, poor CTF fit resolution (>4 Å), and high astigmatism (≥ 1000 Å) were discarded. Initial particles were picked (5,354,890 particles) and extracted from the dose-weighted images with box size of 384 px (Fourier-cropped to 96 px) using CryoSPARC blob picker with dimensions of 80-150 Å and local maxima of 1000 picks. The best particles were aligned and sorted using three rounds of CryoSPARC 2D classification (n = 50) to generate 2D templates for CryoSPARC Template Picker using an average particle diameter of 120 Å. After template picking, initial particles were re-extracted with a box size of 384 px (Fourier-cropped to 128 px) and re-classified through multiple rounds of 2D classification (n=2). The entire dataset consisted of 15,894 motion-corrected images with 2,579,019 particles. Initial models were obtained using CryoSPARC *ab initio* reconstruction (n = 3) using only a subset of the particles (500,000 particles) to generate initial reference-free maps. The *ab initio* reconstruction generated one map representing the IC (1) and two “junk” particle classes (2,3) that were used to curate all extracted particles within the dataset using heterogeneous refinement (n=3), using maps 1 as a reference and the “junk” classes as decoys for particle sorting **(Figure S4)**. After four rounds of heterogeneous refinement, 1,095,540 particles resembling the IC were refined using CryoSPARC non-uniform 3D refinement for downstream processing. To discard particles without bound RNA, a soft mask was generated around the upstream DNA, downstream DNA, and the RNA-DNA hybrid. This nucleic acid soft mask was used as the focus mask for subsequent 3D classification (simple mode, n=6, 5 Å filter resolution) which generated 6 distinct classes of the IC. The 3D classes containing particles with incomplete or weak RNA-DNA hybrid density were omitted from downstream processing **(Figure S4)**. Visual inspection of the two remaining classes resembling the IC in Chimera were identified as IC8 with mtRNAP adopting a closed fingers subdomain conformation with an incoming ATP positioned in the insertion site. Comparison to a recently published substrate bound EC-TEFM complex (8U8V) confirmed our assessment, as there was a clear separation of density between the chain terminator (3’-deoxy GTP) and the incoming ATP nucleotide. To address if any of these particles began transitioning into the elongation phase, thus requiring TFB2M dissociation, we generated a soft mask around TFB2M. This TFB2M soft mask was used as the focus mask for subsequent 3D classification (simple mode, n=3, 5 Å filter resolution, Force Hard Classification) which generated 2 distinct classes representing IC8 and 1 class representing EC8. A final non-uniform refinement (n=2 extra final passes, minimize per-particle scale) of each class resulted in Structures Three, Four, and Five with the following particle counts and nominal resolutions: IC8 (147,533 particles; 2.71 Å), IC8’ (163,170 particles; 2.65 Å), and TC8 (105,850 particles; 2.69 Å).

### Single Particle Analysis of promoter-originated EC9-TEFM

Workflows for image processing of EC9-TEFM are found in **Figure S9**. The EER exposures were imported and processed in CryoSPARC with EER fractionation set to 60 and the EER upsampling factor set to 1. The movies were converted into image stacks and were motion corrected, gain-normalized, drift-corrected, summed, and dose-weighted using the Patch Motion Correction module (CryoSPARC). The Contrast transfer function (CTF) values of the motion corrected movies were estimated using Patch CTF (CryoSPARC). Micrographs with ice, ethane contamination, poor CTF fit resolution (>3 Å), and high astigmatism (≥ 1000 Å) were discarded. Initial particles were picked and extracted from the dose-weighted images with box size of 384 px (Fourier-cropped to 128 px) using CryoSPARC blob picker with dimensions of 100-160 Å and local maxima of 1000 picks. The best particles were aligned and sorted using six rounds of CryoSPARC 2D classification (n = 50) to generate 2D templates for CryoSPARC Template Picker using an average particle diameter of 160 Å. After template picking, initial particles were re-extracted with a box size of 320 px (Fourier-cropped to 104 px) and re-classified through multiple rounds of 2D classification (n=2). The entire dataset consisted of 2,565 motion-corrected images with 998,913 particles. Initial models were obtained using CryoSPARC *ab initio* reconstruction (n = 3) using only a subset of the particles (500,000 particles) to generate initial reference-free maps. The *ab initio* reconstruction generated one map representing the EC-TEFM (1), one map representing the IC (2), and one “junk” particle class (3) that were used to curate all extracted particles within the dataset using heterogeneous refinement (n=3), using maps 1 and 2 as references and the “junk” class as a decoy for particle sorting **(Figure S9)**. After three rounds of heterogeneous refinement, 163,202 particles representing the EC-TEFM were refined using CryoSPARC non-uniform 3D refinement (Punjani *et al*., 2020) for downstream processing. The remaining 150,027 particles within the class resembling the IC were initially considered and processed with 3D Classification and non-uniform refinement, however, these particles did not yield functional ICs containing RNA or interpretable promoter DNA density and were therefore omitted from further processing. To discard particles without nucleic acid in EC-TEFM, a soft mask was generated around the upstream DNA, downstream DNA, and the RNA-DNA hybrid. This nucleic acid soft mask was used as the focus mask for subsequent 3D classification (simple mode, n=6, 6 Å filter resolution) which generated 6 distinct classes resembling the EC-TEFM. The 3D classes containing particles with weak TEFM or RNA-DNA hybrid density were omitted from downstream processing **(Figure S9)**. Visual inspection of the remaining particles in the class representing the EC-TEFM in Chimera was identified as EC9-TEFM with mtRNAP adopting a closed fingers subdomain conformation with an incoming 3’-deoxy ATP positioned in the insertion site. A final non-uniform refinement (n=2 extra final passes, minimize per-particle scale) of EC9-TEFM resulted in Structure Six, comprising 41,738 particles and achieving a nominal resolution of 3.77 Å.

### Single Particle Analysis of promoter-originated EC13

Workflows for image processing of Complex FOUR are found in **Figure S10**. The EER exposures were imported and processed in CryoSPARC^6^ with EER fractionation set to 60 and the EER upsampling factor set to 1. The movies were converted into image stacks and were motion corrected, gain-normalized, drift-corrected, summed, and dose-weighted using the Patch Motion Correction module (CryoSPARC). The Contrast transfer function (CTF) values of the motion corrected movies were estimated using Patch CTF (CryoSPARC). Micrographs with ice, ethane contamination, poor CTF fit resolution (>4 Å), and high astigmatism (≥ 1000 Å) were discarded. Initial particles were picked and extracted from the dose-weighted images with box size of 384 px (Fourier-cropped to 128 px) using CryoSPARC blob picker with dimensions of 80-140 Å and local maxima of 1000 picks. The best particles were aligned and sorted using three rounds of CryoSPARC 2D classification (n = 50). The entire dataset consisted of 11,693 motion-corrected images with 2,525,605 particles. Initial models were obtained using CryoSPARC *ab initio* reconstruction (n = 3) using only a subset of the particles (500,000 particles) to generate initial reference-free maps. The *ab initio* reconstruction generated one map representing the EC (1), one map representing the IC (2), and one “junk” particle class (3) that were used to curate all extracted particles within the dataset using heterogeneous refinement (n=3), using maps 1 and 2 as references and the “junk” class as a decoy for particle sorting **(Figure S10)**. After four rounds of heterogeneous refinement, 583,723 particles representing the EC were refined using CryoSPARC non-uniform 3D refinement for downstream processing. The remaining 547,493 particles within the class resembling the IC were initially considered and processed with 3D Classification and non-uniform refinement, however, these particles did not yield functional ICs containing RNA or interpretable promoter DNA density and were therefore omitted from further processing. To discard particles without bound RNA in the EC, a soft mask was generated around the upstream DNA, downstream DNA, and the RNA-DNA hybrid. This nucleic acid soft mask was used as the focus mask for subsequent 3D classification (simple mode, n=6, 5 Å filter resolution) which generated 6 distinct classes resembling the EC. The 3D classes containing particles with apo-mtRNAP or incomplete or weak RNA-DNA hybrid density were omitted from downstream processing **(Figure S10)**. Visual inspection of the remaining particles in the class representing the EC in Chimera (Pettersen et al., 2021) was identified as EC13 with mtRNAP adopting a closed fingers subdomain conformation with an incoming 3’-deoxy CTP positioned in the insertion site. A final non-uniform refinement (n=2 extra final passes, minimize per-particle scale) of EC13 resulted in Structure Seven, comprising 43,172 particles and achieving a nominal resolution of 2.74 Å.

### Post-Processing of Cryo-EM Maps

Post-processing of all of the cryo-EM density maps were performed by DeepEMhancer (Sanchez-Garcia et al., 2021) using the highRes or tightTarget models. The reported resolutions of the cryo-EM maps are based on the Fourier shell correlation (FSC) 0.143 criterion (Scheres and Chen, 2012). Local resolution calculations were generated using CryoSPARC Local Resolution Estimation and displayed in ChimeraX (Pettersen *et al*., 2021) **(Figures S2,5,6,11).** The angular distribution of particle orientations, conical FSC (cFSC) and cFSC Area Ratio (cFAR) summary plots, and weighted distribution of viewing directions were assessed by the CryoSPARC Orientation Diagnostics module and were calculated for all structures **(Figures S2,5,6,11)**.

### Model Building and Structure Refinement

Initial models for the initiation complex and elongation complexes were derived from the Protein Data Bank (PDB) using PDB IDs 6ERP^4^ and 8U8V^3^. The models were manually docked into the respective cryo-EM maps using USCF Chimera^9^. DNA B-form and RNA A-form restraints derived from Coot v.0.9.8.2 (Casanal et al., 2020) were used to fit the polynucleotide chains in the non-template DNA, template DNA, and RNA primer strands. The ATP substrates for IC8, IC8’, TC8, and EC9-TEFM were obtained from the Coot monomer library and were fit using the Jiggle-Fit Ligand module in Coot. The 3’-deoxy CTP substrate for EC13 was generated by importing a SMILES string for 3’-deoxy CTP (PubChem CID 153054) using Coot’s Ligand Builder, and restraints were generated from refinement within Coot. Model building was assisted with iterative rounds of real-space structure refinement using PHENIX 1.21.1 (Afonine et al., 2018) using Ramachandran and secondary structure restraints. Models were inspected and modified in Coot and the refinement process was repeated iteratively to obtain the final structures. Comprehensive model validation was carried out with PHENIX (Adams et al., 2010) and the PDB validation server (https://validate-rcsb-2.wwpdb.org/) and is summarized in **(Tables S1-3)**. Figures and movies were generated with PyMOL (Schrödinger, LLC) and ChimeraX.

## Notes

### Competing Interest Statement

The authors have declared no competing interest.

