## Supplemental Figures and Tables for "Structural Basis for Promoter Recognition and Transcription Factor Binding and Release in Human Mitochondria"

The file includes Supplemental Figures S1-S13 and Tables S1-S3

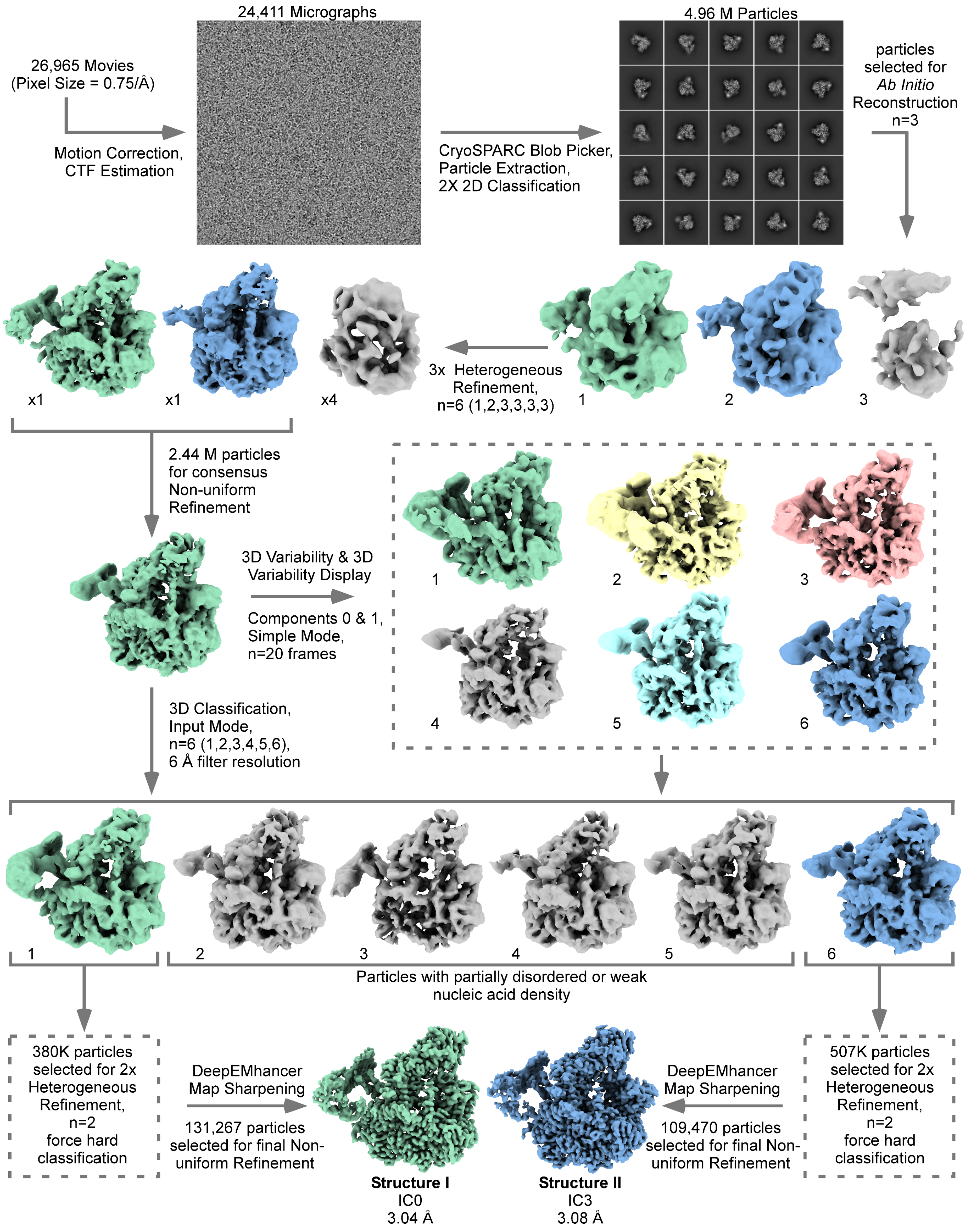


**Figure S1. Related to Figure 1. | Cryo-EM data processing workflow for early-stage human mitochondrial ICs.** Flow chart of cryo-EM single-particle analysis with a representative micrograph, 2D class images, and maps of the IC particles.

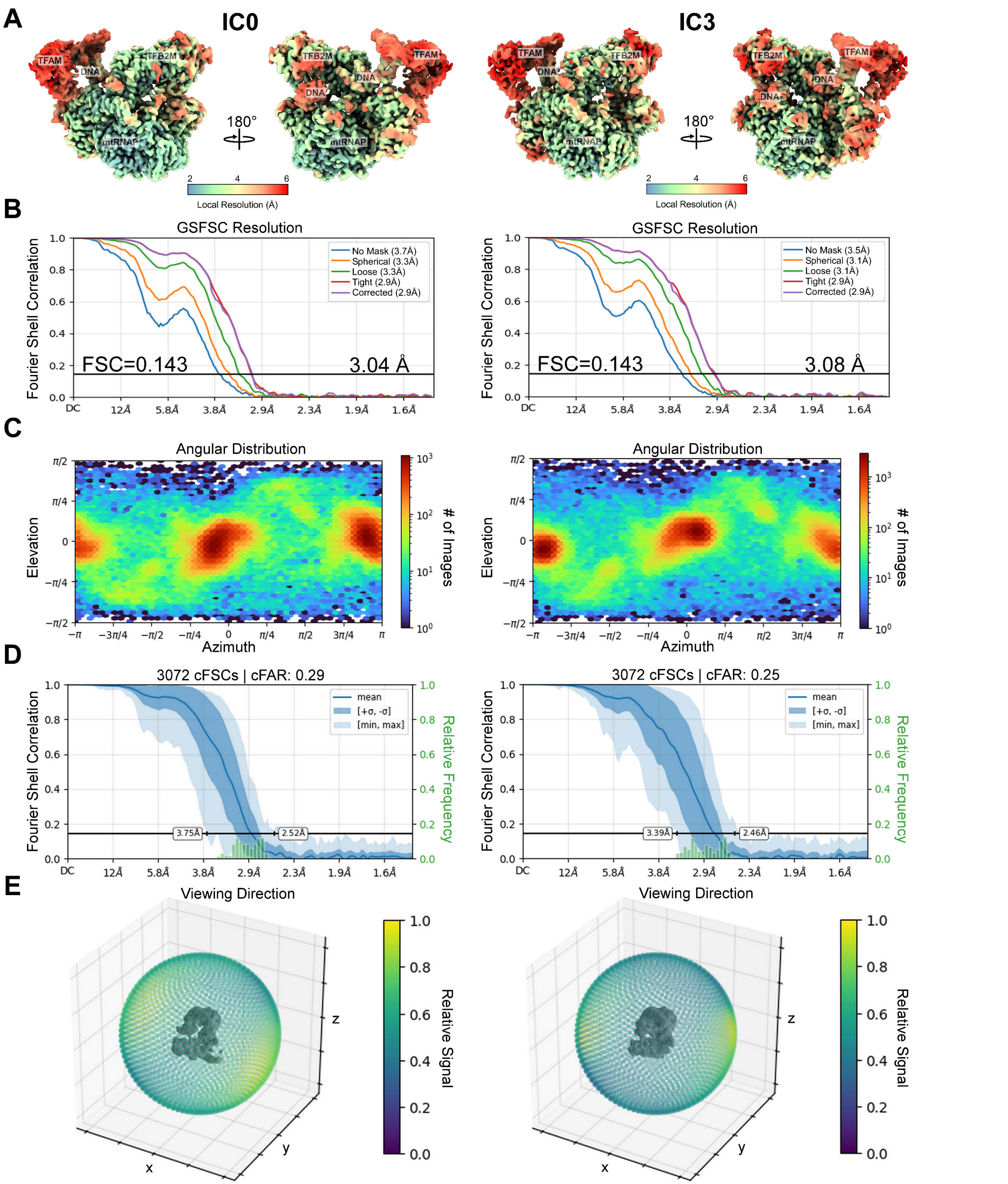


**Figure S2. Related to Figure 1. | Cryo-EM quality assessment of IC0 and IC3 structures. (A)** Local resolution estimation of IC0 (*left*) and IC3 (*right*) using CryoSPARC and surface colored in ChimeraX. **(B)** Gold Standard FSC curves from CryoSPARC (FSC threshold 0.143). **(C)** Distribution of viewing angles and orientations for the cryo-EM density maps. **(D)** Orientation diagnostics showing the conical FSC area ratio (cFAR) and **(E)** relative signal visualized by a 3D-colored viewing sphere encompassed by lowpass filtered maps.

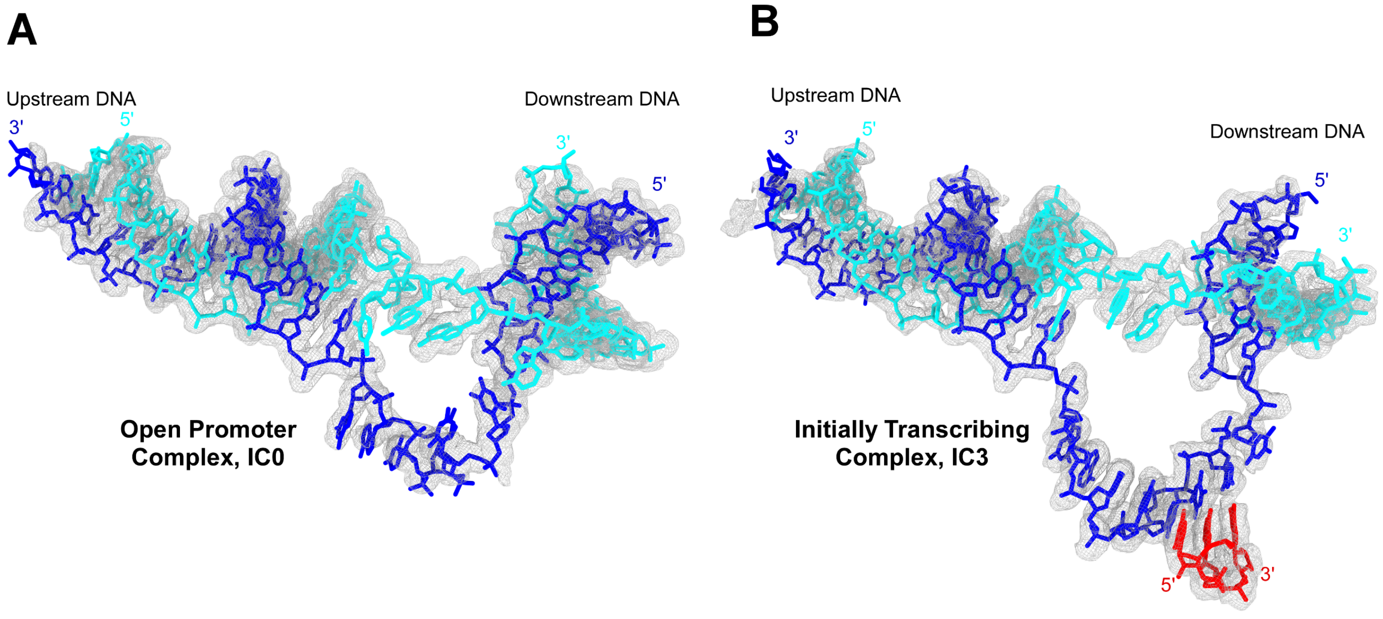


**Figure S3. Related to Figure 1. | Cryo-EM density maps of nucleic acids in the early-stage ICs.** Cryo-EM densities of the DNA transcription bubble in the open promoter complex, IC0 **(A)**, and the initially transcribing complex, IC3 **(B)**. The NT strand, T strand, and RNA are colored in cyan, blue, and red, respectively.

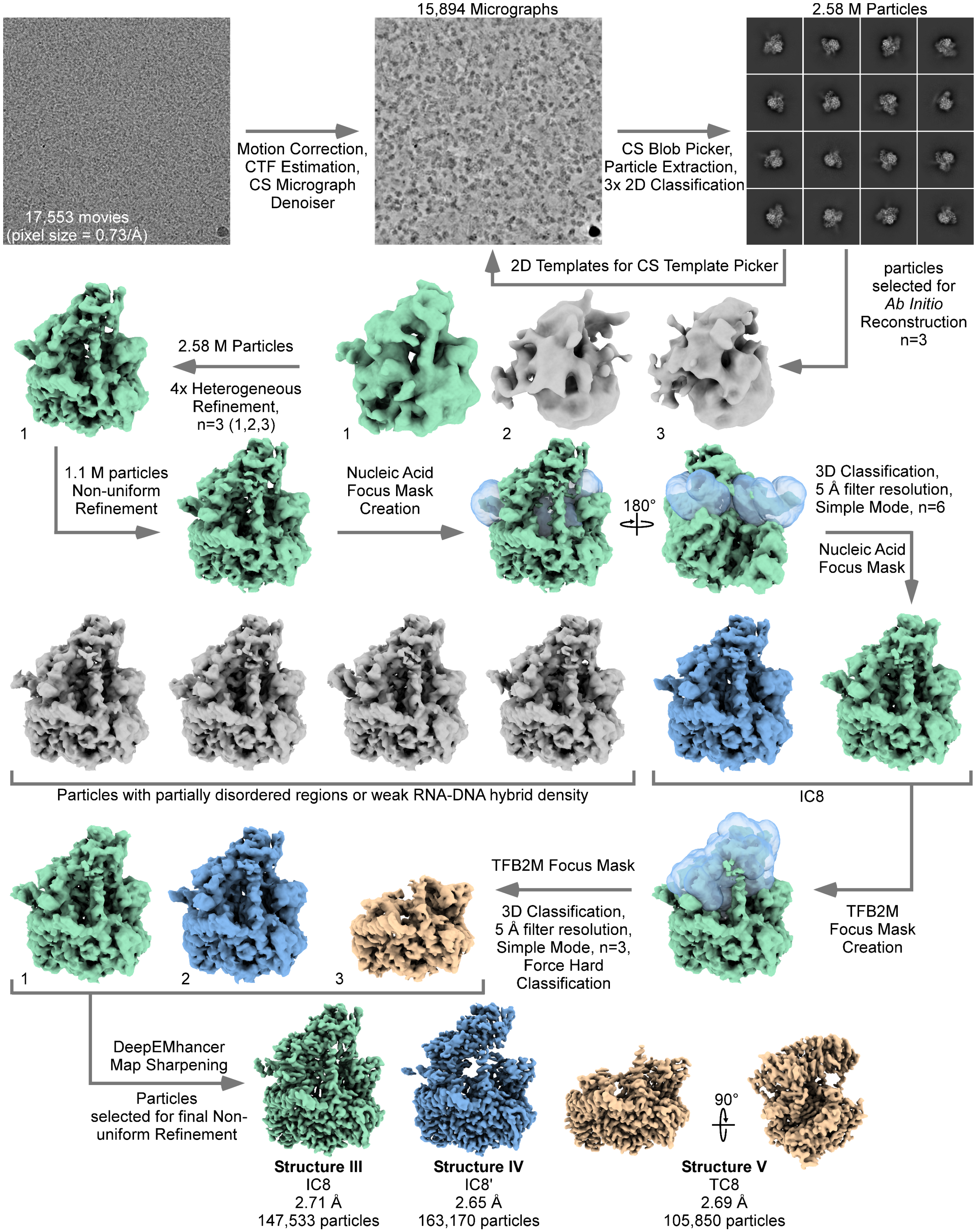


**Figure S4. Related to Figure 4. | Cryo-EM data processing workflow for the late-stage mitochondrial ICs.** Flow chart of cryo-EM single-particle analysis with representative micrographs, 2D class images, and maps of the IC particles

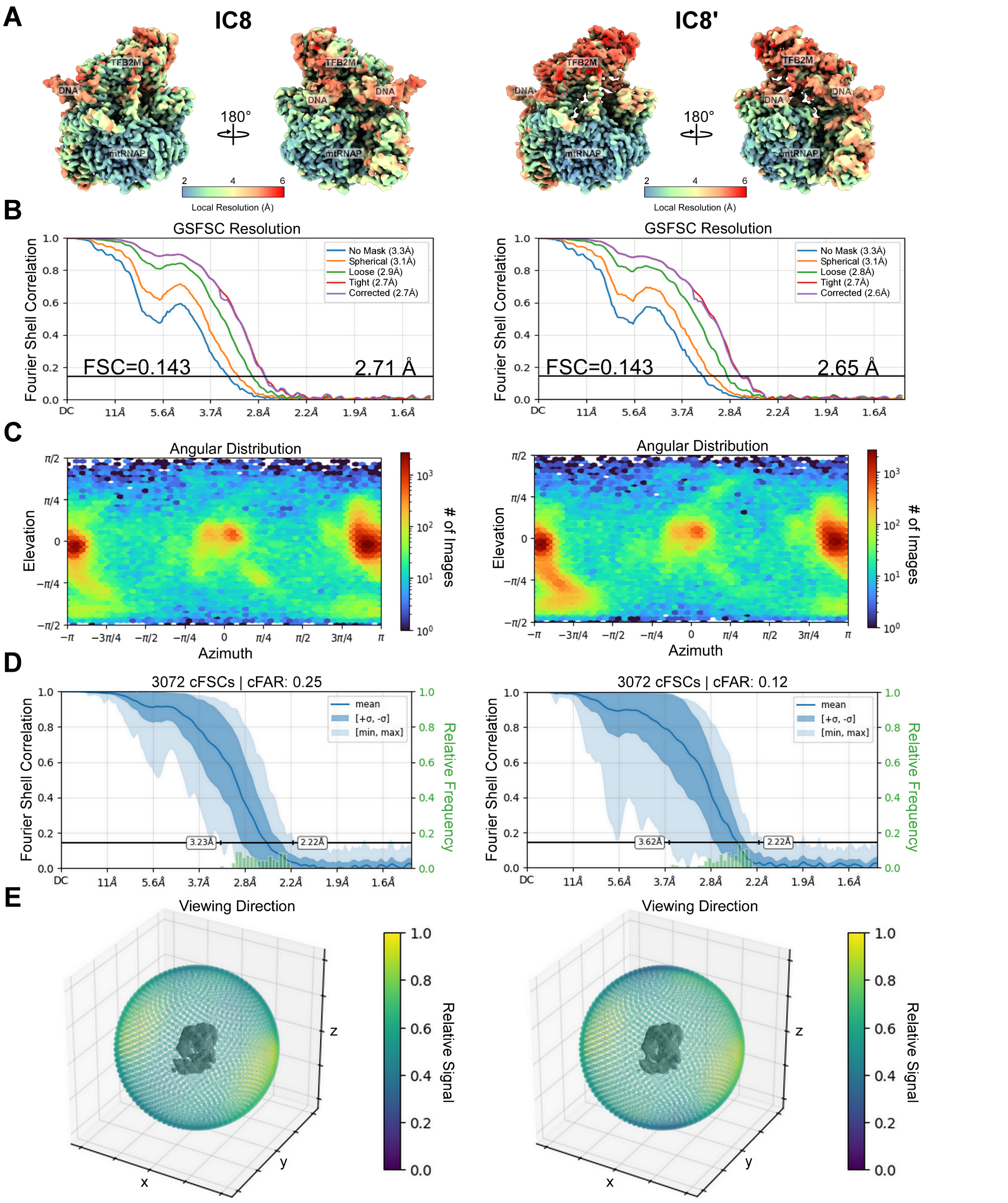


**Figure S5. Related to Figure 4. | Cryo-EM quality assessment of IC8 and IC8' structures. (A)** Local resolution estimation of IC8 (*left*) and IC8' (*right*) using CryoSPARC and surface colored in ChimeraX. **(B)** Gold Standard FSC curves from CryoSPARC (FSC threshold 0.143). **(C)** Distribution of viewing angles and orientations for the cryo-EM density maps. **(D)** Orientation diagnostics showing the conical FSC area ratio (cFAR) and **(E)** relative signal visualized by a 3D-colored viewing sphere encompassed by lowpass filtered maps.

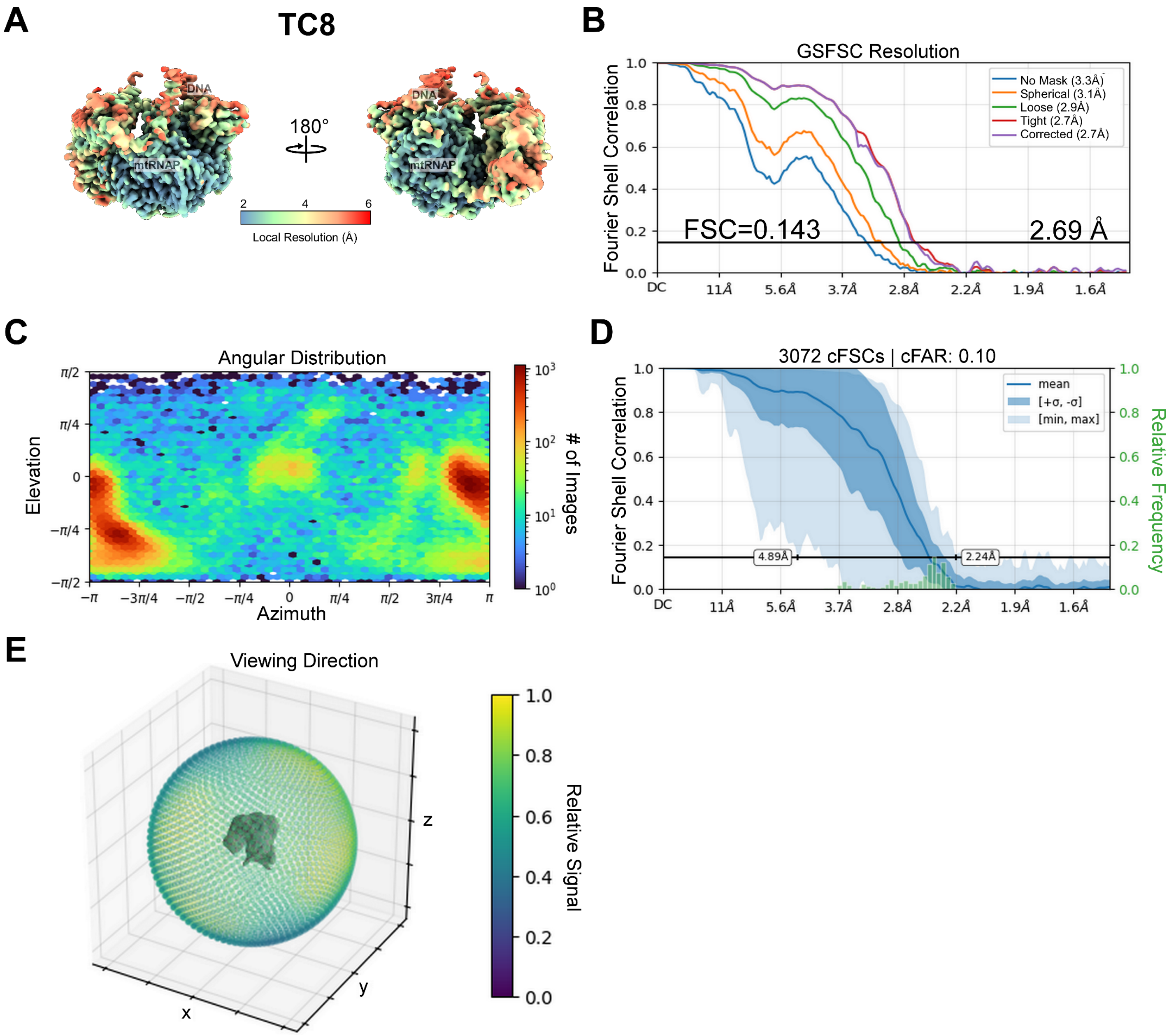


**Figure S6. Related to Figure 4. | Cryo-EM quality assessment of the TC8 structure. (A)** Local resolution estimation of TC8 using CryoSPARC and surface colored in ChimeraX. **(B)** Gold Standard FSC curves from CryoSPARC (FSC threshold 0.143). **(C)** Distribution of viewing angles and orientations for the cryo-EM density maps. **(D)** Orientation diagnostics showing the conical FSC area ratio (cFAR) and **(E)** relative signal visualized by a 3D-colored viewing sphere encompassed by lowpass filtered maps.

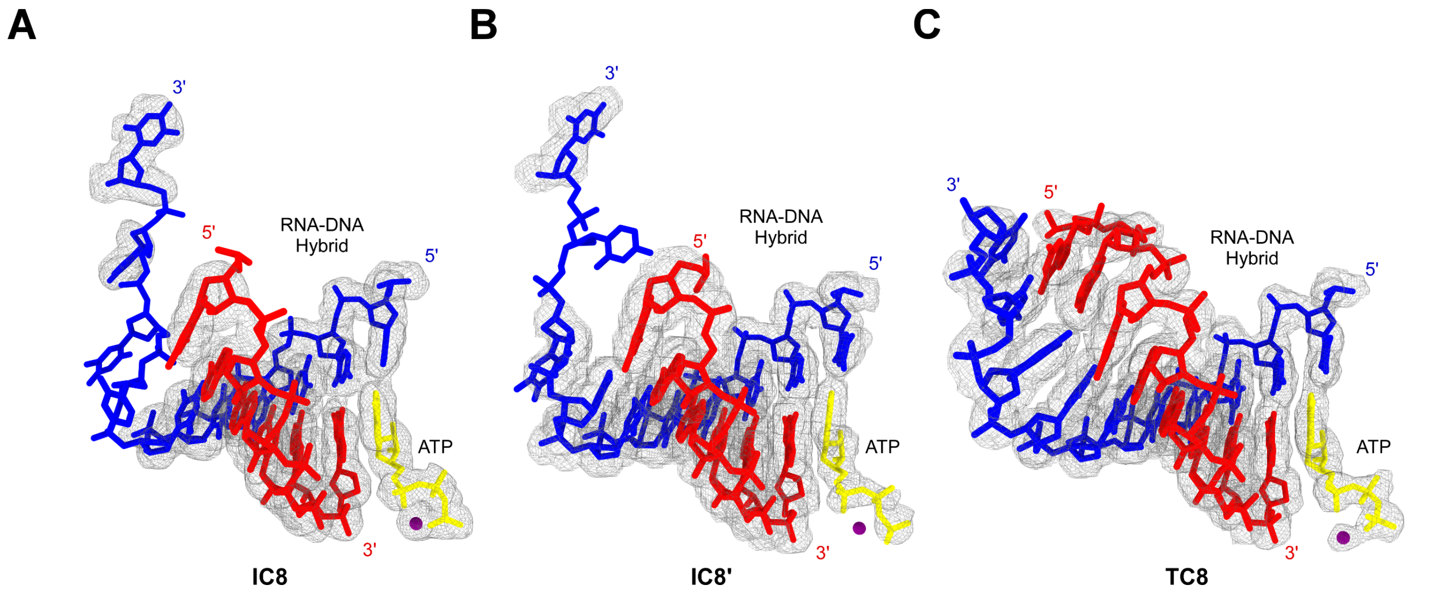


**Figure S7. | Cryo-EM density maps of nucleic acids in the late-stage ICs.** Cryo-EM densities of the RNA-DNA hybrids in IC8 **(A)**, IC8' **(B)**, and TC8 **(C)**. The T strand, RNA, substrate ATP, and magnesium ion are colored in blue, red, yellow, and purple, respectively.

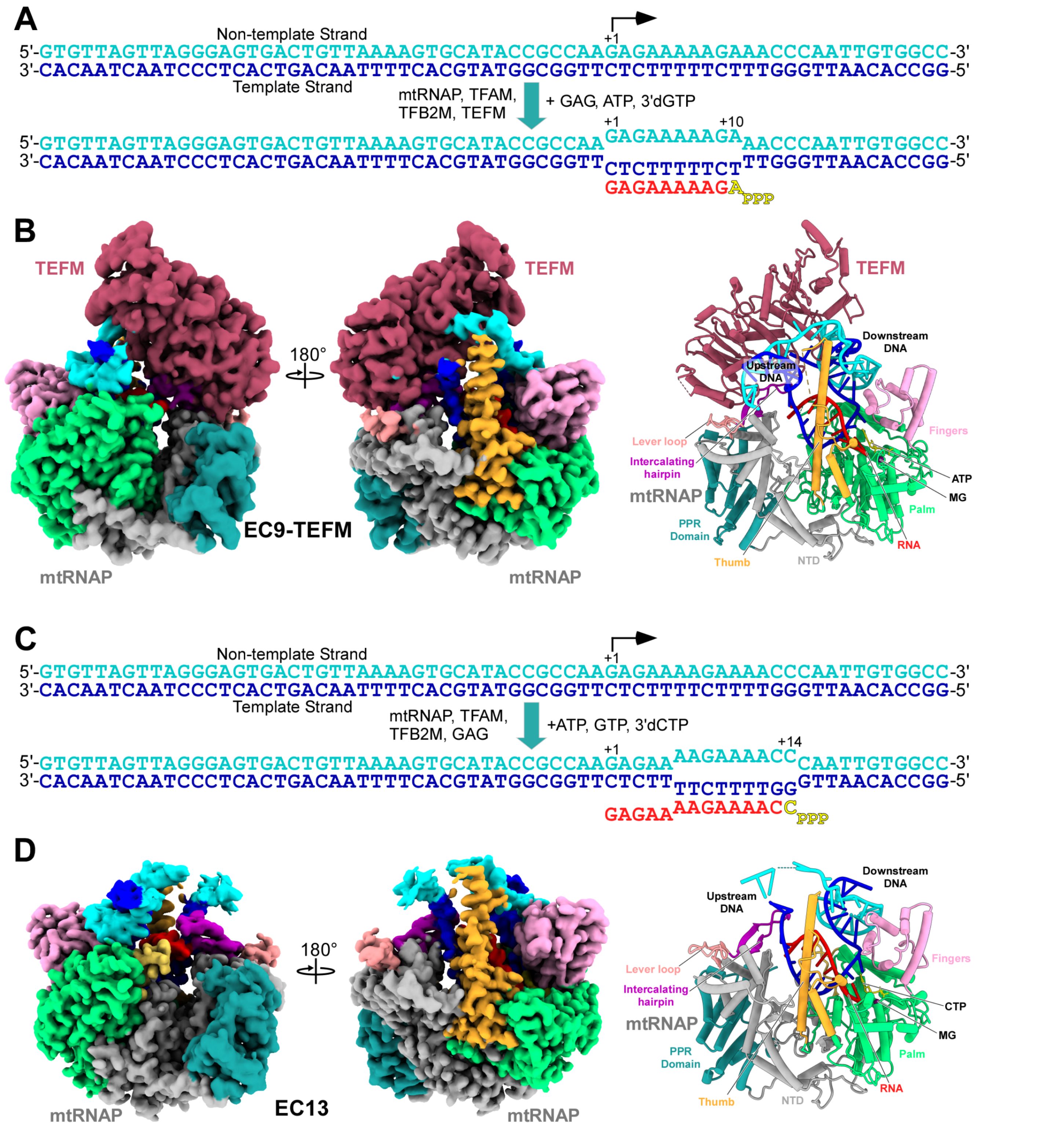


**Figure S8. Related to Figure 6. | Structure of the promoter-originated transcription elongation complexes.**

**(A)** Design of a fully double-stranded promoter template used to generate EC9. The TSS is indicated at +1. The NT DNA strand is shown in cyan, TS – in blue, and RNA – in red. The incoming ATP in the insertion site is depicted in yellow. **(B)** The cryo-EM density and the structure of EC9-TEFM. **(C)** Design of a fully double-stranded promoter template used to generate EC13. **(D)** The cryo-EM density and structure of EC13.

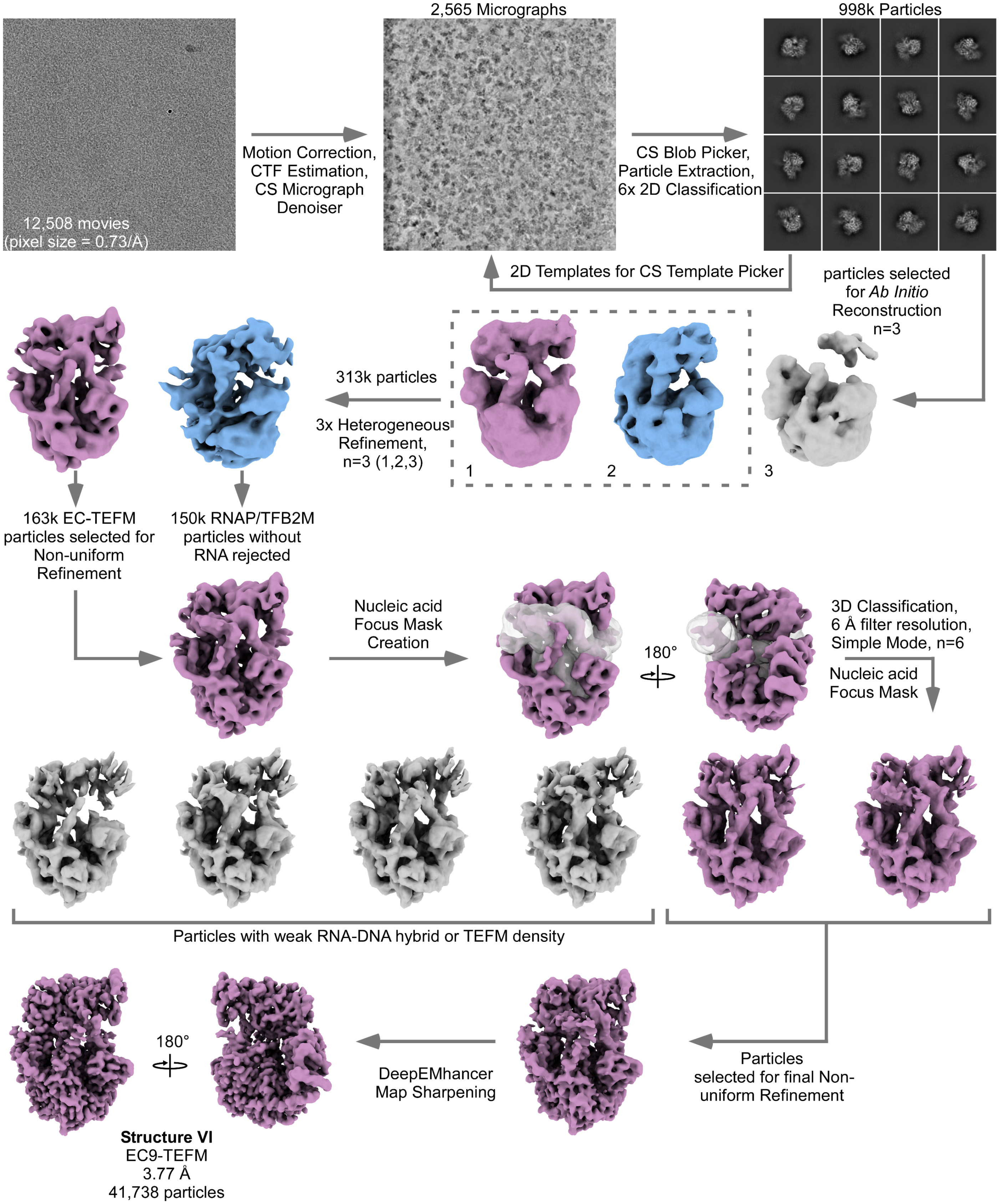


**Figure S9. Related to Figure 6. | Cryo-EM data processing workflow for EC9-TEFM.** Flow chart of cryo-EM single-particle analysis with representative micrographs, 2D class images, and maps of the EC-TEFM particles.

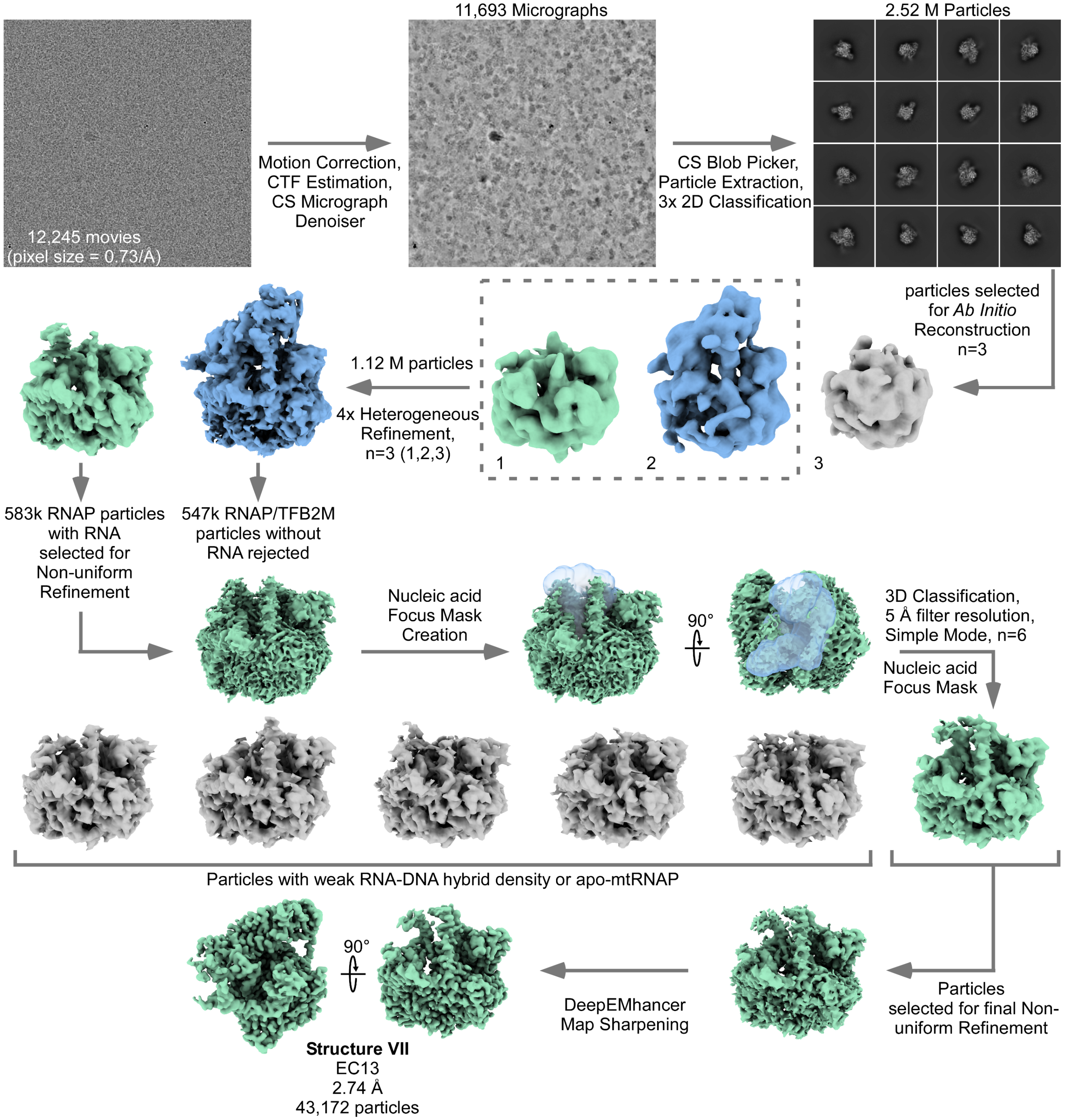


**Figure S10. Related to Figure 6. | Cryo-EM data processing workflow for EC13.** Flow chart of cryo-EM single-particle analysis with representative micrographs, 2D class images, and maps of the particles.

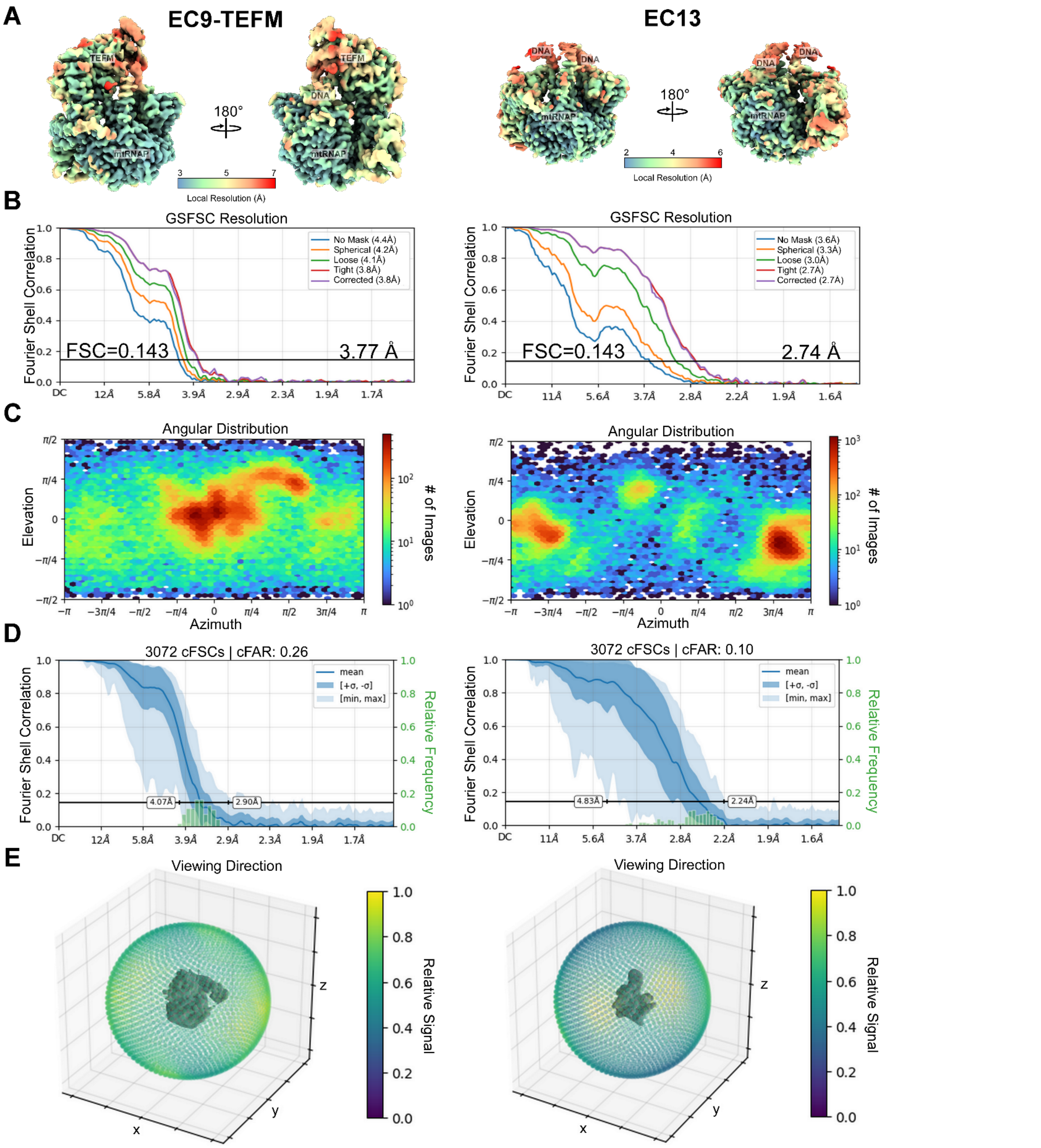


**Figure S11. Related to Figure 6. | Cryo-EM quality assessment of EC9-TEFM and EC13 structures. (A)** Local resolution estimation of EC9-TEFM (*left*) and EC13 (*right*) using CryoSPARC and surface colored in ChimeraX. **(B)** Gold Standard FSC curves from CryoSPARC (FSC threshold 0.143). **(C)** Distribution of viewing angles and orientations for the cryo-EM density maps. **(D)** Orientation diagnostics showing the conical FSC area ratio (cFAR) and **(E)** relative signal visualized by a 3D-colored viewing sphere encompassed by lowpass filtered maps.

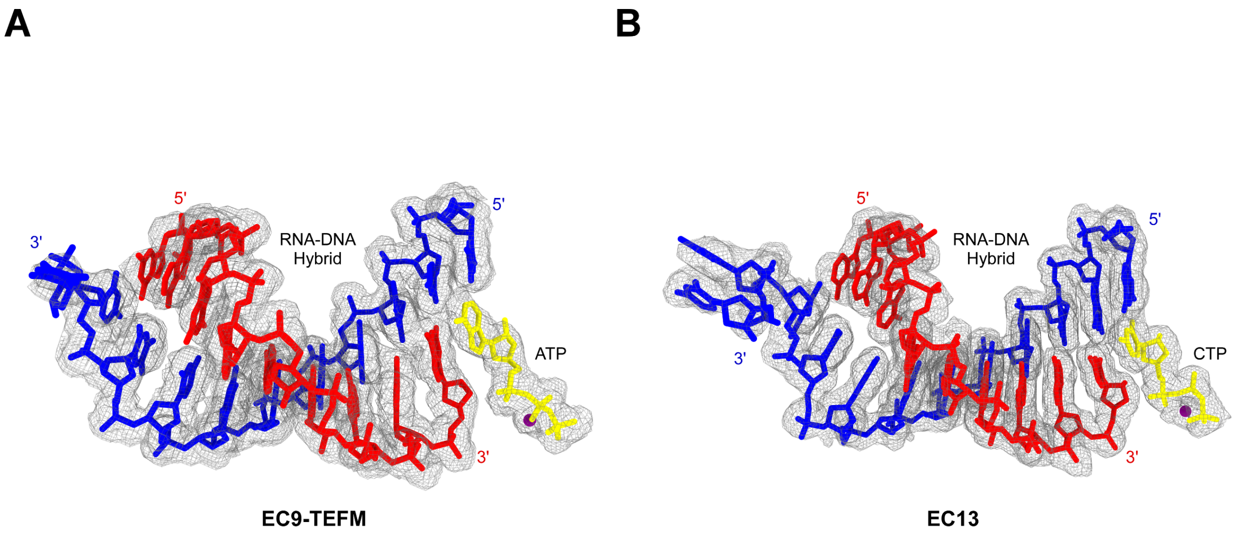


**Figure S12. Related to Figure 6. | Cryo-EM density maps of the RNA-DNA duplex in EC9-TEFM (A) and EC13 (B).** The T strand, RNA, substrate ATP, and magnesium ion are colored in blue, red, yellow, and purple, respectively.

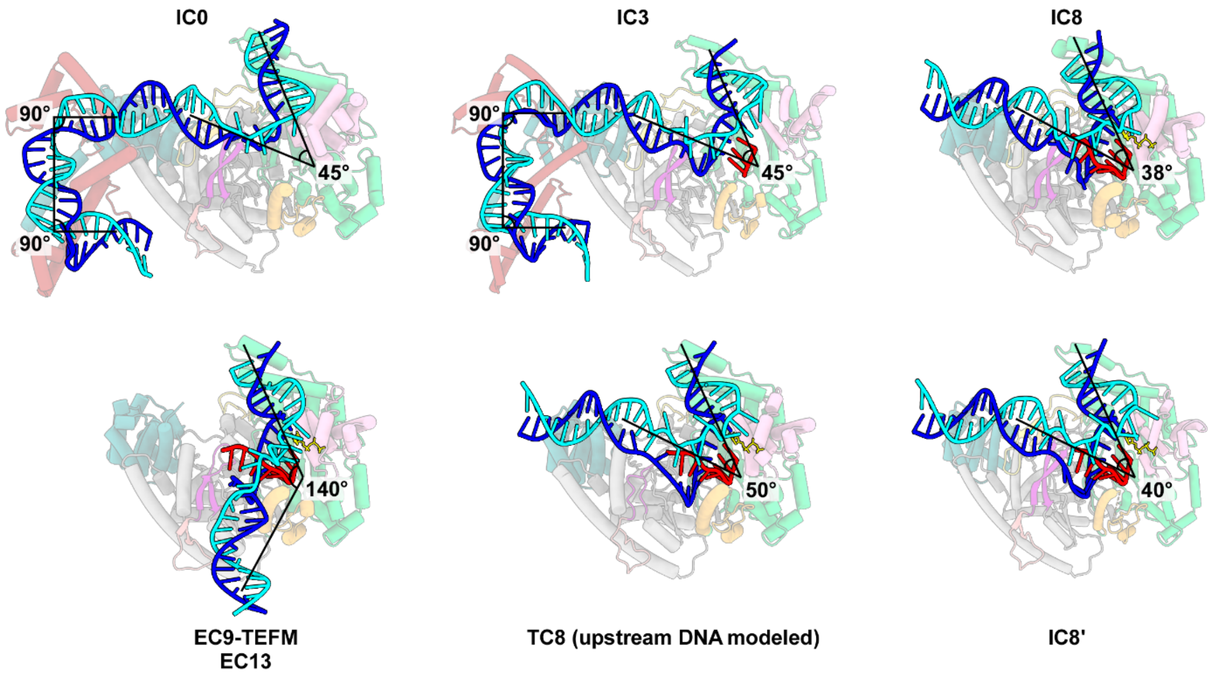


**Figure S13. | Related to Figure 7. Changes in DNA topology in mitochondrial transcription complexes during the transition from initiation to elongation.** Ribbon representation of DNA bending observed in the IC and EC structures. Structures contain mtRNAP and TFAM in the background and DNA in the foreground. Transcription factors TFB2M and TEFM were omitted for clarity. Helices are depicted as cylinders. The approximate angles between DNA duplexes are indicated.

**Table S1 | Cryo-EM data collection, refinement, and validation statistics of Early-Stage ICs**

| **Dataset** | **COMPLEX ONE (IC LSP50 -3/+1 Template + GAG)** | |
|---|---|---|
| **Data Collection** | **Structure One: IC0** | **Structure Two: IC3** |
| Electron Microscope | Titan Krios | Titan Krios |
| Voltage (kV) | 300 | 300 |
| Detector | Falcon 4i | Falcon 4i |
| Camera mode | Counting | Counting |
| Magnification | 165,000x | 165,000x |
| Total electron dose (e- $Å^{-2}$) | 47.69 | 47.69 |
| Defocus range (- µm) | 0.3-2.5 | 0.3-2.5 |
| Pixel Size (Å) | 0.7511 | 0.7511 |
| Grid type | 300 mesh Holey Carbon (~5 nm gold layer) nanowire | 300 mesh Holey Carbon (~5 nm gold layer) nanowire |
| Software | Leginon | Leginon |
| Dose rate (e- $Å^{-2}$ $s^{-1}$) | 7.95 | 7.95 |
| Frames (no.) | 60 | 60 |
| Energy Filter | Selectris 10 eV | Selectris 10 eV |
| Micrographs collected (no.) | 26,965 | 26,965 |
| **Image Processing** | | |
| SPA Software | CryoSPARC 4.5.3 | CryoSPARC 4.5.3 |
| Symmetry imposed | C1 | C1 |
| Initial particles picked (no.) | 7,863,657 | 7,863,657 |
| Particles picked after 2D classification (no.) | 4,957,265 | 4,957,265 |
| Particles picked after het. refinement (no.) | 2,441,562 | 2,441,562 |
| Particles picked after 3D classification (no.) | 380,094 | 506,855 |
| Final particles picked (no.) | 131,267 | 109,470 |
| Map Resolution (Å) | 3.04 | 3.08 |
| FSC Threshold | 0.143 | 0.143 |
| **Refinement** | | |
| Initial model used (PDB) | 6ERP | 6ERP |
| Map sharpening B factor ($Å^2$) | 75.9 | 73.9 |
| Model composition | | |
| *Nonhydrogen atoms* | 14759 | 14835 |
| *Protein residues* | 1578 | 1578 |
| *Nucleotides* | 100 | 103 |
| *Water* | 0 | 0 |
| *Ligands* | 0 | 0 |

| | | |
|---|---|---|
| B factors ($Å^2$) (min/max/mean) | | |
| *Protein* | 28.02/521.01/326.42 | 29.67/98.79/87.69 |
| *Nucleotide* | 20.00/612.97/336.76 | 20.00/612.97/242.46 |
| *Ligand* | --/--/-- | --/--/-- |
| R.m.s. deviations | | |
| *Bond lengths (Å) (# > 4σ)* | 0.004 (0) | 0.004 (0) |
| *Bond angles (°) (# > 4σ)* | 0.820 (1) | 0.808 (7) |
| Validation | | |
| *MolProbity score* | 1.45 | 1.56 |
| *Clash Score* | 5.14 | 5.32 |
| Poor rotamers (%) | 0.00 | 0.00 |
| Ramachandran Plot | | |
| *Preferred (%)* | 97.00 | 96.04 |
| *Allowed (%)* | 3.00 | 3.96 |
| *Outliers (%)* | 0.00 | 0.00 |

**Table S2 | Cryo-EM data collection, refinement, and validation statistics of late-stage ICs**

| **Dataset** | **COMPLEX TWO (IC LSP66 Template + GAG, ATP, dGTP)** | | |
|---|---|---|---|
| **Data Collection** | **Structure Three: IC8** | **Structure Four: IC8'** | **Structure Five: TC8** |
| Electron Microscope | Titan Krios | Titan Krios | Titan Krios |
| Voltage (kV) | 300 | 300 | 300 |
| Detector | Falcon 4i | Falcon 4i | Falcon 4i |
| Camera mode | Counting | Counting | Counting |
| Magnification | 165,000x | 165,000x | 165,000x |
| Total electron dose (e- $Å^{-2}$) | 45.44 | 45.44 | 45.44 |
| Defocus range (- μm) | 0.4-3.2 | 0.4-3.2 | 0.4-3.2 |
| Pixel Size (Å) | 0.7304 | 0.7304 | 0.7304 |
| Grid type | 300 mesh Holey gold Ultra-AuFoil 1.2/1.3 | 300 mesh Holey gold Ultra-AuFoil 1.2/1.3 | 300 mesh Holey gold Ultra-AuFoil 1.2/1.3 |
| Software | Leginon | Leginon | Leginon |
| Dose rate (e- $Å^{-2}$ $s^{-1}$) | 7.57 | 7.57 | 7.57 |
| Frames (no.) | 60 | 60 | 60 |
| Energy Filter | Selectris 10 eV | Selectris 10 eV | Selectris 10 eV |
| Micrographs collected (no.) | 17,559 | 17,559 | 17,559 |
| **Image Processing** | | | |
| SPA Software | CryoSPARC 4.6.0 | CryoSPARC 4.6.0 | CryoSPARC 4.6.0 |
| Symmetry imposed | C1 | C1 | C1 |
| Initial particles picked (no.) | 5,354,890 | 5,354,890 | 5,354,890 |
| Particles picked after 2D classification (no.) | 2,579,019 | 2,579,019 | 2,579,019 |
| Particles picked after het. refinement (no.) | 1,095,540 | 1,095,540 | 1,095,540 |
| Particles picked after 3D classification (no.) | 416,553 | 416,553 | 416,553 |
| Final particles picked (no.) | 147,533 | 163,170 | 105,850 |
| Map Resolution (Å) | 2.71 | 2.65 | 2.81 |
| FSC Threshold | 0.143 | 0.143 | 0.143 |
| **Refinement** | | | |
| Initial model used (PDB) | 6ERP | 6ERP | 8U8V |
| Map sharpening B factor ($Å^2$) | 63.0 | 61.0 | 59.1 |
| Model composition | | | |
| *Nonhydrogen atoms* | 11856 | 11789 | 8500 |
| *Protein residues* | 1306 | 1298 | 971 |
| *Nucleotides* | 66 | 66 | 35 |
| *Water* | 0 | 0 | 0 |
| *Ligands* | ATP:1, MG:1 | ATP:1, MG:1 | ATP:1, MG:1 |
| B factors ($Å^2$) (min/max/mean) | | | |

| | | | |
|---|---|---|---|
| *Protein* | 30.00/463.78/110.82 | 30.00/463.78/111.28 | 30.00/463.78/120.57 |
| *Nucleotide* | 20.00/75.21/37.21 | 20.00/75.21/36.95 | 20.00/79.49/32.36 |
| *Ligand* | 20.00/20.00/20.00 | 20.00/20.00/20.00 | 20.00/20.00/20.00 |
| R.m.s. deviations | | | |
| *Bond lengths (Å) (# > 4σ)* | 0.005 (1) | 0.005 (0) | 0.006 (0) |
| *Bond angles (º) (# > 4σ)* | 0.877 (8) | 0.934 (7) | 0.998 (9) |
| Validation | | | |
| *MolProbity score* | 1.50 | 1.58 | 1.53 |
| *Clash Score* | 5.69 | 4.89 | 5.14 |
| Poor rotamers (%) | 0.00 | 0.00 | 0.00 |
| Ramachandran Plot | | | |
| *Preferred (%)* | 96.84 | 95.35 | 96.16 |
| *Allowed (%)* | 3.16 | 4.65 | 3.84 |
| *Outliers (%)* | 0.00 | 0.00 | 0.00 |

**Table S3 | Cryo-EM data collection, refinement, and validation statistics of ECs**

| Dataset | COMPLEX THREE (IC LSP66 Template + GAG, ATP, dGTP) | COMPLEX FOUR (IC LSP66 Template + GAG, ATP, GTP, dCTP) |
|---|---|---|
| **Data Collection** | **Structure Six: EC9-TEFM** | **Structure Seven: EC13** |
| Electron Microscope | Titan Krios | Titan Krios |
| Voltage (kV) | 300 | 300 |
| Detector | Falcon 4i | Falcon 4i |
| Camera mode | Counting | Counting |
| Magnification | 165,000x | 165,000x |
| Total electron dose (e- $Å^{-2}$) | 46.52 | 45.44 |
| Defocus range (- µm) | 0.4-2.7 | 0.3-2.2 |
| Pixel Size (Å) | 0.7304 | 0.7304 |
| Grid type | 300 mesh Holey gold Ultra-AuFoil 1.2/1.3 | 300 mesh Holey gold Ultra-AuFoil 1.2/1.3 |
| Software | Leginon | Leginon |
| Dose rate (e- $Å^{-2}$ $s^{-1}$) | 7.75 | 7.42 |
| Frames (no.) | 60 | 60 |
| Energy Filter | Selectris 10 eV | Selectris 10 eV |
| Micrographs collected (no.) | 12,508 | 12,245 |
| **Image Processing** | | |
| SPA Software | CryoSPARC 4.6.0 | CryoSPARC 4.6.0 |
| Symmetry imposed | C1 | C1 |
| Initial particles picked (no.) | 1,574,006 | 4,696,418 |
| Particles picked after 2D classification (no.) | 998,913 | 2,525,605 |
| Particles picked after het. refinement (no.) | 313,229 | 1,133,162 |
| Particles picked after 3D classification (no.) | 163,202 | 583,723 |
| Final particles picked (no.) | 41,738 | 43,172 |
| Map Resolution (Å) | 3.77 | 2.62 |
| FSC Threshold | 0.143 | 0.143 |
| **Refinement** | | |
| Initial model used (PDB) | 8U8V | 8U8V |
| Map sharpening B factor ($Å^2$) | 113.7 | 44.9 |
| Model composition | | |
| *Nonhydrogen atoms* | 12076 | 8742 |
| *Protein residues* | 1383 | 977 |
| *Nucleotides* | 69 | 44 |
| *Water* | 0 | 0 |
| *Ligands* | ATP:1, MG:1 | dCTP:1, MG:1 |
| B factors ($Å^2$) (min/max/mean) | | |

| | | |
|---|---|---|
| *Protein* | 30.00/286.49/152.13 | 30.00/463.78/121.79 |
| *Nucleotide* | 20.00/319.98/119.63 | 20.00/79.49/31.15 |
| *Ligand* | 20.00/30.00/20.31 | 20.00/20.00/20.00 |
| R.m.s. deviations | | |
| *Bond lengths (Å) (# > 4σ)* | 0.004 (0) | 0.005 (0) |
| *Bond angles (º) (# > 4σ)* | 0.867 (7) | 0.938 (3) |
| Validation | | |
| *MolProbity score* | 1.62 | 1.46 |
| *Clash Score* | 6.42 | 4.73 |
| Poor rotamers (%) | 0.00 | 0.00 |
| Ramachandran Plot | | |
| *Preferred (%)* | 96.13 | 96.60 |
| *Allowed (%)* | 3.87 | 3.40 |
| *Outliers (%)* | 0.00 | 0.00 |
